# Mechanisms by Which Small Molecule Inhibitors Arrest Sec14 Phosphatidylinositol Transfer Protein Activity

**DOI:** 10.1101/2022.08.01.502361

**Authors:** Xiao-Ru Chen, Lokendra Poudel, Zebin Hong, Philipp Johnen, Sachin S. Katti, Ashutosh Tripathi, Aaron H. Nile, Savana M. Green, Gabriel Schaaf, Fulvia Bono, Vytas A. Bankaitis, Tatyana I. Igumenova

## Abstract

Phosphatidylinositol transfer proteins (PITPs) promote phosphoinositide signaling by enhancing phosphatidylinositol (PtdIns) 4-OH kinase activities in producing signaling pools of PtdIns-4-phosphate. As such, PITPs are key regulators of lipid signaling in eukaryotic cells. While the PITP phospholipid exchange cycle is the engine that stimulates PtdIns 4-OH kinase activity, the protein and lipid dynamics associated with this critical process are not understood. Herein, we use an integrative structural approach that takes advantage of small molecule inhibitors (SMIs) directed against the major yeast PITP (Sec14) to gain new insights into the mechanics of the Sec14 phospholipid exchange cycle from the perspective of protein, phospholipid and SMI dynamics. Moreover, as Sec14 has emerged as an attractive target for next-generation antifungal drugs, the structures of Sec14 bound to SMIs of four different chemotypes reported in this study provide critical information required for structure-based design of next-generation lead compounds that target Sec14 PITPs of virulent fungi.

## Introduction

Phosphatidylinositol (PtdIns) is the metabolic precursor of phosphoinositides (PIPs). The seven known PIPs are chemically distinguished by their differential phosphorylation patterns at the 3- OH, 4-OH, and/or 5-OH positions on the PtdIns inositol ring. This limited cohort of PIPs, along with their metabolic by-products (e.g. diacylglycerol and soluble inositol-phosphates), regulate an impressively broad set of intracellular activities. As such, PIPs represent the foundation of a major pathway for intracellular signaling in all eukaryotic cells, and the 4-OH phosphorylated PIPs PtdIns-4-phosphate (PtdIns-4-P) and PtdIns-4,5-bisphosphate are essential for viability at the single cell level across the *Eukaryota* (Di Paolo and De Camilli, 2006; Strahl and Thorner, 2007; Balla, 2013; Dickson and Hille, 2019).

While contemporary research efforts focus primarily on the lipid kinases and phosphatases that produce and consume PIPs, respectively, key questions regarding how PIP production is coordinated with the larger metabolome remain. It is in those contexts that PtdIns transfer proteins (PITPs) emerge as novel regulators of PIP signaling – particularly of PtdIns-4- P signaling. PITPs fall into two structurally unrelated Sec14-like and START (StAR-related lipid transfer)-like PITP families defined by their respective founding members – the yeast Sec14 and the mammalian PITPα (Grabon et al., 2019). The biological importance of Sec14 and START PITPs is abundantly evident in both uni- and multi-cellular organisms. PITP deficiencies in individual PITPs of either family are associated with striking phenotypes in fungi, plants, insects, and vertebrates (Bankaitis et al., 1989; Milligan et al., 1997; Vincent et al., 2005; Giansanti et al., 2006; Ile et al., 2010, Huang et al., 2016; Koe et al., 2018) -- including mammals (Hamilton et al., 1999; Alb et al., 2003; Xie et al., 2018).

Current models envision the *S. cerevisiae* Sec14 and its homologs as PtdIns- presentation modules that stimulate PtdIns-4-P synthesis. Sec14-mediated PtdIns presentation to PtdIns 4-OH kinases is proposed to be driven by a lipid exchange cycle by which Sec14 exchanges PtdCho for PtdIns on the surface of yeast trans-Golgi membranes. This cycle is suggested to be initiated by membrane recruitment of a closed Sec14 conformer from the cytoplasm where bound lipid is shielded from the aqueous environment (Fig. 1A step 1). Sec14 then is proposed to undergo a conformational transition to the open form where lipid exchange between bilayer and Sec14 lipid-binding cavity can occur (Fig. 1A step 2) until a turn of the exchange cycle results in disengagement of a closed lipid-bound Sec14 conformer from the membrane surface (Fig. 1A, step 3). The process of lipid exchange renders PtdIns a superior substrate for PtdIns 4-OH kinases that are otherwise intrinsically poor interfacial enzymes incapable of overcoming the actions of antagonists of PtdIns-4-P signaling (e.g. Sac1 PtdIns-4- P phosphatase and Kes1/Osh proteins; Schaaf et al., 2008; Bankaitis et al., 2010). The functional partnership between Sec14 and PtdIns 4-OH kinase translates into a PtdCho-based metabolic input that activates membrane trafficking from the yeast Golgi system via activation of effectors that promote downstream PtdIns-4-P signaling events (Fig. 1A, step 4).

**Figure 1.**
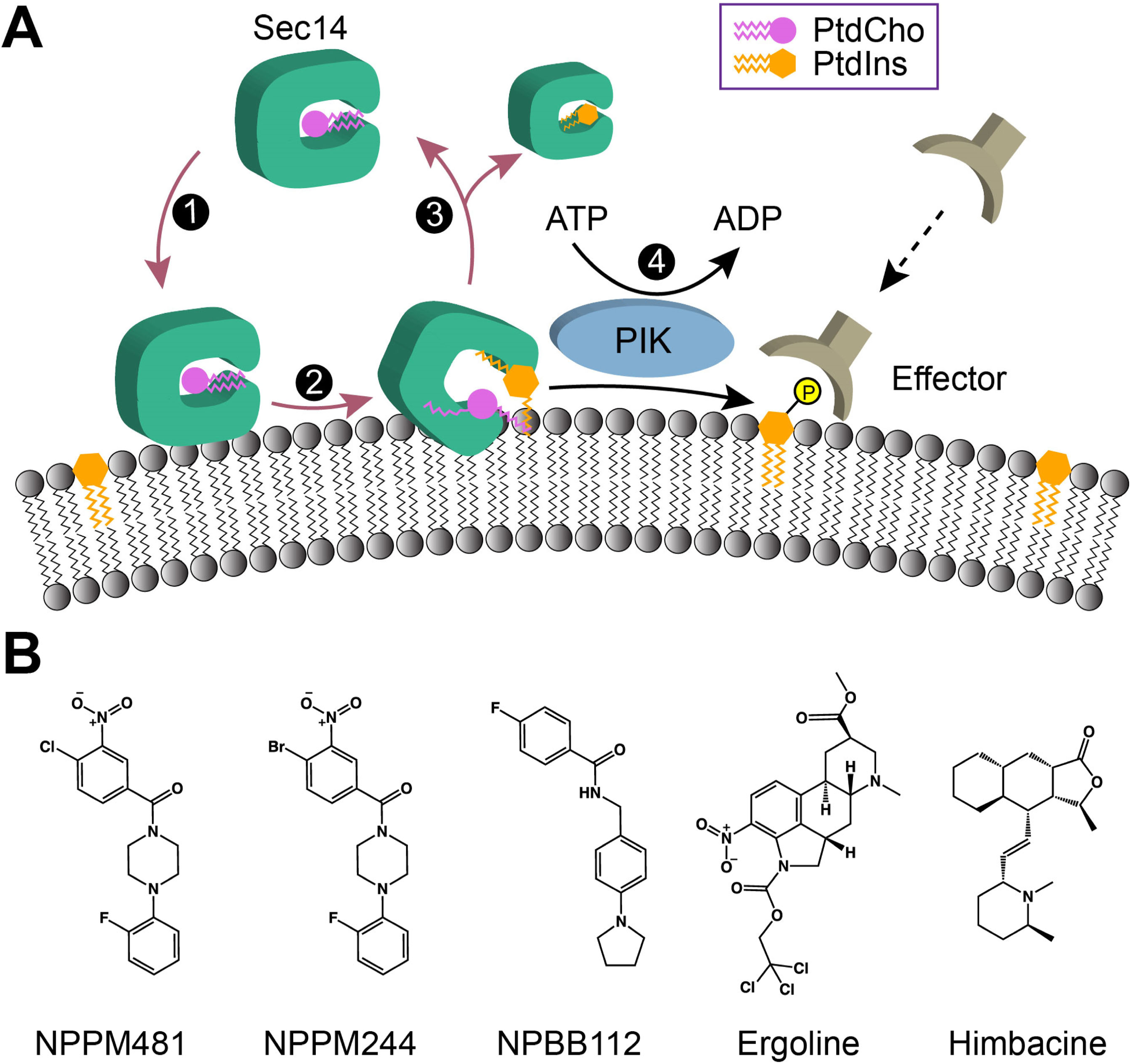
The Sec14 phospholipid exchange cycle stimulates PtdIns-4-P production by PtdIns 4-OH kinases. **(A)** PtdIns presentation model for Sec14-mediated potentiation of PtdIns-4-P signaling. PtdIns 4-OH kinases are biologically insufficient interfacial enzymes that cannot produce sufficient PtdIns-4-P to overcome the action of antagonists of PtdIns-4- P signaling. Recruitment of Sec14 to trans-Golgi network membranes (1) activates a PtdIns/PtdCho exchange cycle (2). PtdIns/PtdCho exchange ultimately results in Sec14 disengagement from membranes (3), but execution of the lipid exchange cycle(s) renders PtdIns a superior substrate for PtdIns 4-OH kinase (4). In this manner, Sec14 stimulates lipid kinase activity such that sufficient PtdIns-4-P is generated to produce a biologically actionable signal. **(B)** Sec14-directed small molecule inhibitors of four distinct chemotypes. Chemical structures of the 4-chloro- and 4-bromo-3-nitrophenyl)(4-(2- methoxyphenyl)piperazin-1-yl)methanones NPPM481 and NPPM244, respectively, the 4- fluoro-N-[4-(1-pyrrolidinyl)benzyl]benzamide) NPBB112, the ergoline NGx04, and the piperidine alkaloid natural product himbacine are shown. These SMIs inhibit the Sec14 phospholipid exchange cycle.

Small molecule inhibitors (SMIs) of five distinct chemotypes have been validated to target yeast Sec14 with exquisite specificity (Nile, 2014; Nile et al., 2014, Pries et al., 2018). SMIs that specifically target Sec14 hold dual promise. First, Sec14-directed SMIs hold potential as tool compounds for dissecting the lipid exchange cycle mechanism. This exchange cycle is central to PtdIns-4-P signaling, but the protein and lipid dynamics associated with it are not at all understood. Second, fungal Sec14s are emerging as promising new targets for development of urgently needed next-generation anti-mycotics (Nile et al., 2014; Khan et al., 2021: Bankaitis et al., 2022). Sec14-directed SMIs serve as valuable lead compounds for such efforts.

Herein, we investigate the underlying mechanisms of inhibition by Sec14-directed SMIs of four distinct chemotypes (Fig. 1B). Using an integrative structural approach consisting of protein crystallography, molecular dynamics simulations and fluorine-19 Nuclear Magnetic Resonance (^19^F NMR) spectroscopy, we provide a comprehensive structural and quantitative description of Sec14-directed SMI interactions with their target. These findings lend new insights into Sec14-mediated lipid exchange and provide ‘proof-of-concept’ that Sec14-directed SMIs are useful tool compounds for dissecting the lipid-exchange cycle. Moreover, the structural information described herein provides key information for drug discovery efforts aimed at designing next-generation anti-mycotics that target Sec14 PITPs of virulent fungi.

## Results

### Crystallization of Sec14 in complex with SMIs of four distinct chemotypes

Sec14-directed SMIs exhibit a range of potencies as measured by inhibition of PtdIns transfer activity in vitro: NPPM244 (IC_50_ = 0.1μM) > NPPM481 (IC_50_ = 0.2 μM) > NPBB112 (IC_50_ = 1.0 μM) > himbacine (IC_50_ = 1.2 μM) (Nile, 2014; Nile et al., 2014). Although the NGxO4 potency in that assay has not been determined, the relative NGxO4 IC_50_ in yeast growth assays (∼10μM) suggests it is a less potent Sec14 inhibitor (Fillipuzzi et al., 2016). Complexes of Sec14 bound to NPPM481, NPPM244, NPBB112, NGx04 (subsequently referred to as ergoline), and himbacine were crystallized and the structures determined at 2.3 Å, 2.1 Å, 2.7 Å, 2.3 Å, and 1.8 Å resolution, respectively. Superposition of the Sec14::SMI complexes with the ‘open’ conformation of apo Sec14 (PDB entry 1AUA) show that the backbone conformations are very similar, with pairwise overall backbone RMSD values ranging from 0.6 to 1.2 Å. Per-residue RMSD values mapped onto the corresponding protein structures reveal that the largest deviations from the reference structure involve the α-10 helical gate element (Gly_210_ to Ile_242_; Fig. 2A). However, as this structural element is involved in crystal contact in all five cases, we do not ascribe larger functional significance to these deviations. Additionally, in the himbacine complex, helical regions α5 - α7 and the N-terminal segment of helix α9, have per-residue RMSD values that exceed 2 Å.

**Figure 2.**
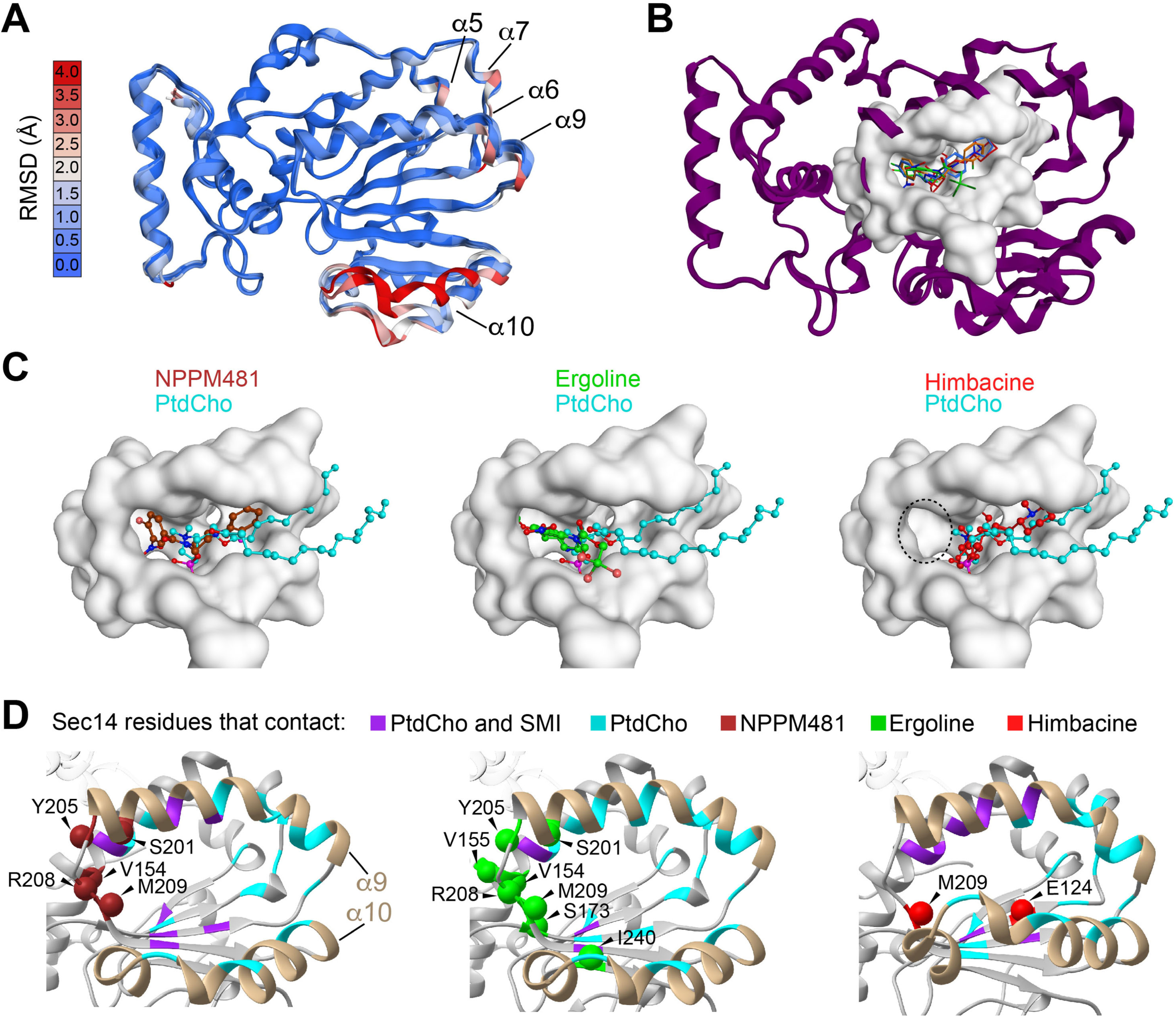
Small molecule inhibitors occupy the PtdCho-binding space. **(A)** Per-residue RMSD values, color-coded and mapped onto each of the Sec14::SMI crystal structures. The apo Sec14 that crystallizes as an open conformer (1AUA) is used as reference. α-Helices that include regions with significant RMSD values are labeled. Color-coded scale bar is at left. **(B)** Overlay of the binding poses of the five SMIs in the context of the Sec14 fold are shown. Highlighted in space-fill render is a twenty-one amino acid SMI-binding envelope that accommodates SMIs of all four chemotypes. The NPPM481, NPBB112, ergoline, and himbacine backbones are in maroon, blue, green, and red, respectively. Cl and F atoms are shown in pink and magenta, respectively. NPPM244 is not represented in the overlay as it superposes precisely onto the NPPM481 pose. **(C)** An expanded view of the NPPM481, ergoline and himbacine poses in the common binding envelope, as indicated. The PtdCho pose (cyan) is provided for reference in each case, and is extrapolated from the Sfh1::PtdCho structure reported by Schaaf et al (2008). The subpocket occupied by NPPMs/NPBB/ergoline but not himbacine is marked with a dotted circle. **(D)** Ribbon representations of open Sec14 lipid binding cavities of the Sec14::NPPM481 (left), Sec14::ergoline (center) and Sec14::himbacine complexes (right) are shown. The α9 and α10 helices that gate entry to the cavity are labeled. Beads mark residues that contact SMI only and the identities of those residues are shown. Regions that engage both SMI and PtdCho are colored in purple in the ribbon diagram, whereas those that engage only PtdCho are shown in cyan.

Although the SMIs represent four distinct chemotypes, all five compounds are sequestered in what we define as a common envelope of twenty-one amino acids that form an amphiphilic subregion deep within the Sec14 lipid-binding cavity (Fig. 2B). In that regard, all SMI binding poses overlap the Sec14 binding space dedicated to accommodation of the natural ligand PtdCho -- specifically its headgroup, glycerol backbone and the proximal acyl chain regions (Fig 2C). As NPPM481 and NPPM244 differ only in the identity of the A-ring halide substituent, these SMIs share essentially identical binding modes within the Sec14 lipid-binding cavity, and the NPBB112 binding mode also deviates in only minor respects (Fig. S1A-C). The himbacine molecule is set much shallower in the collective SMI binding environment than are the NPPMs/NPBB/ergoline SMIs – thereby leaving vacant the subpocket occupied by those SMIs (Fig. 2C, dotted circle).

Considering only hydrophobic and H-bond interactions, eleven of the twenty-one Sec14 residues that form the SMI binding environment interact directly with PtdCho (Fig. S1A-F). The unique sets of Sec14 residues that interact with individual SMIs are highlighted using color-coding and bead representation in Fig. 2D and are compared to the cohort of Sec14 residues that interact with the natural ligand PtdCho. Ergoline and himbacine engage in the most and the least number of unique interactions, respectively. To emphasize the comparative analysis of Sec14-SMI interactions, we organize the subsequent sections according to the identities of the SMI chemical moieties and their positions within the lipid-binding pocket.

### NPPM/NPBB A-ring and ergoline carboxylic ester group (R1) occupy a common hydrophobic pocket adjacent to the PtdCho headgroup binding region

The A-rings of NPPM481, NPPM244, NPBB112 and the ergoline R1 group occupy a compact and mostly hydrophobic pocket that resides adjacent to (but does not overlap) the PtdCho headgroup-binding motif, whereas himbacine does not (Figs. 3A,B,C). This pocket is formed by residues Tyr_111_, Tyr_151_, Val_154_, Val_155_, Ser_173_, Tyr_205_ and Arg_208_. The aryl-halide A-ring of NPPM481 is sandwiched by π-π stacking interactions with Tyr151 and van der Waals contacts with Arg208 and Tyr205 (Fig. 3A,D). The Cl atom is wedged into a hydrophobic cavity lined by the Val_154_ and Val_155_ methyl groups, the Arg_208_ methylenes, and the Tyr_151_ aromatic ring. We find no evidence for halogen-bond interactions of NPPM481 or NPPM244 with Ser_173_ as had previously been suggested by docking simulations (Nile et al., 2014). Instead, the halide (Cl or Br) engages in ‘edge-on’ halogen-π interactions (Imai et al., 2008) with the Tyr_151_ aromatic ring in both structures -- as illustrated for NPPM481 in Fig. 3D. The position of the NPBB112 A-ring corresponds closely to the A-ring position of Sec14::NPPM481 (Fig. S2A), but the small radius and low polarizability of the A-ring F substituent precludes the formation of hydrophobic and halogen-π interactions in the pocket. Previous SAR analyses identified the nature and the position of the A-ring halide as essential features of an active Sec14-targeted NPPM (Nile et al., 2014). The NPPM structures rationalize the SAR data.

**Figure 3.**
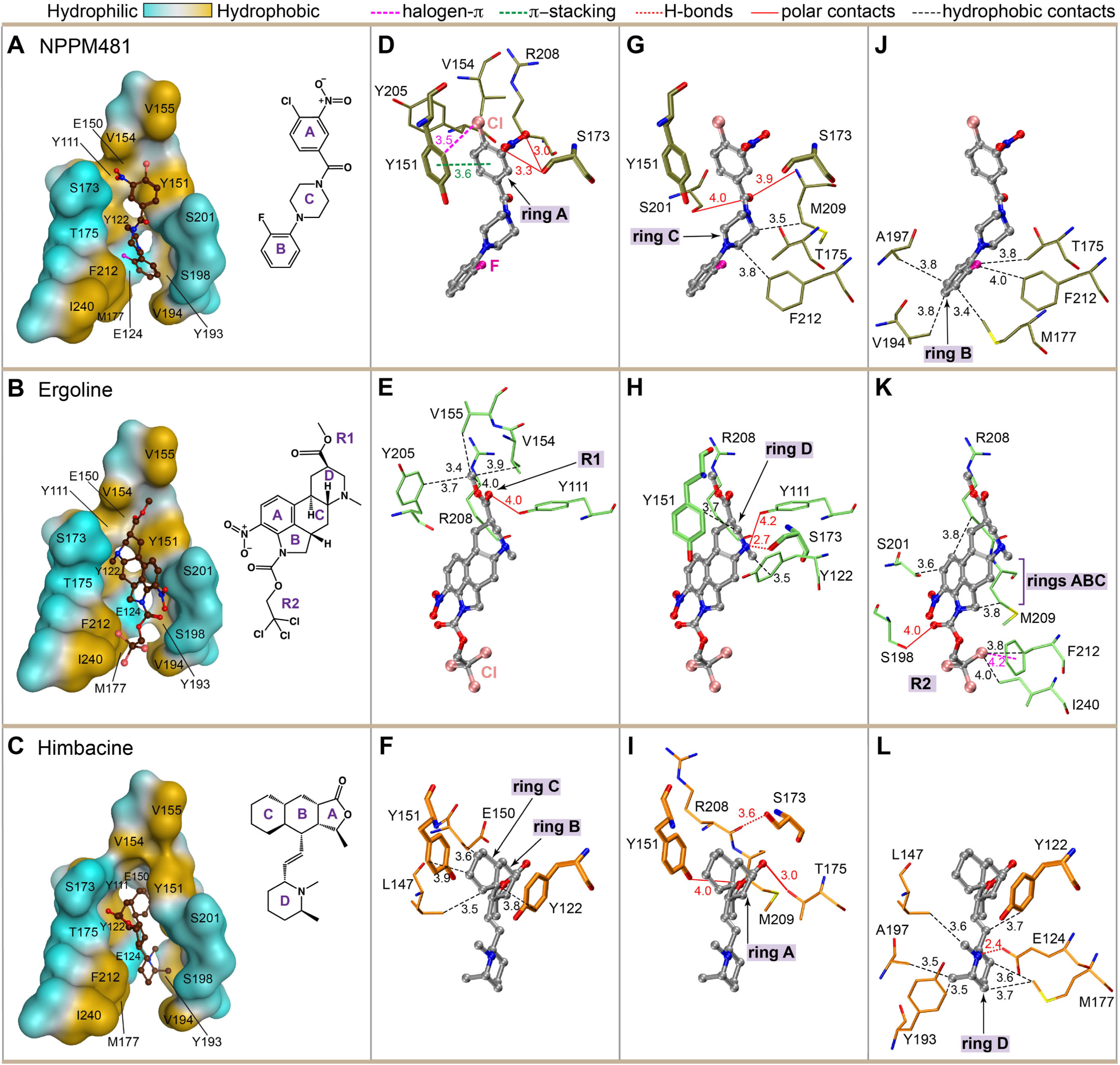
High resolution descriptions of NPPM481, ergoline and himbacine binding modes. Poses of NPPM481, ergoline and himbacine are shown in cut-away views of the common twenty-one amino acid binding envelope (Panels **A**,**B**,**C**, respectively). Amino acids depicted in the cut-away are labeled. Tyr_205_, Arg_208_, and Met_209_ are omitted to not obstruct the view. Leu_147_ and Ala_197_ are not visible in this orientation. The orientation of the envelope is rotated 90 degrees clockwise relative the orientation shown in Figure 2. The chemical structures of the SMIs are presented in the same orientation as these appear in the panels of the corresponding rows. Panels **D**,**G**,**J** depict the binding modes for the NPPM481 A-ring, C-ring and linker, and B-ring, respectively. In **(D)**, Tyr_111_ is omitted to not obstruct the view of other interactions. The OH group of the Tyr_111_ sidechain forms a hydrogen bond with the nitro group of the NPPM481 A-ring. Panels **E**,**H**,**K** detail Sec14 interactions with the ergoline carboxylic ester, fused ring system and tricholoroethyl-carboxylic ester constituents. Panels **F**,**I**,**L** describe interactions between Sec14 and the himbacine B and C rings, the A-ring, and the D-ring, respectively. The intra-protein H-bond between Ser_173_ and Arg_208_ is shown in (I) to emphasize the polar character of the region. Atom color scheme: carbon (gray), oxygen (red), nitrogen (dark blue), chlorine (pink) and fluorine (magenta). Interaction key: halide-π interactions (dashed magenta lines), π-π interactions (green dashed lines), hydrophobic interactions (dashed black lines), polar interactions (solid red lines), hydrogen bonds (dashed red lines). Corresponding distances are indicated in Å.

As is the case for the NPPM481/NPPM244 halides, the ergoline carboxylic ester methyl group represents a major node of interaction with Sec14 as it engages in van der Waals contacts with Val_154_, Val_155_, Arg_208_, and Tyr_205_ (Fig. 3E). With respect to polar interactions, a key residue is Tyr_111_ whose sidechain serves as the H-bond donor to the oxygens of the NPPM481 nitro group (Fig. S1C), and forms a polar contact with the ergoline carbonyl oxygen (Fig. 3E). Ser_173_ additionally contributes to the hydrophilic environment of the nitro group of NPPM481.

### Binding modes for the NPPM C-ring and ergoline and himbacine fused ring systems

The NPPM481linker and C-ring, and the ergoline and himbacine fused ring systems, occupy a common region of the SMI binding envelope. This region provides a restricted binding space with significant hydrophilic character, and it overlaps the PtdCho headgroup and glycerol backbone-binding space. The NPPM481 linker and C-ring exhibit polar character and are accommodated in a narrow hydrophilic ‘neck’ flanked by Tyr_151_ and Thr_175_ hydroxyl groups. The Ser_173_ and Ser_201_ sidechain hydroxyls and the Met_209_ backbone amide contribute to the polar nature of this restricted binding space (Fig. 3G). The NPBB112 carboxamide is similarly accommodated in this hydrophilic ‘neck’ with the Ser_173_ hydroxyl and the Met_209_ backbone amide on one side and the Tyr_151_ hydroxyl on the other. Binding of the NPBB112 C-ring is stabilized by van der Waals contacts with Met_209_ and Phe_212_ (Fig. S1A).

The ergoline fused ring system is comprised of four rings designated ABCD (Fig. 3B). It is convenient to consider the interactions of the D-ring and the ABC rings with Sec14 separately. The D-ring is situated in a hydrophilic neck where its tertiary nitrogen is flanked by an H-bond with Ser_173_, a polar contact with Tyr_111_ and a potential methyl-π interaction with Tyr_122_ (Fig. 3H). The D-ring hydrophobic interactions involve van der Waals contacts with the aromatic ring carbons of Tyr_151_. The ABC-rings are bracketed by an extended hydrophobic methylene spine formed by the sidechains of Arg_208_ and Met_209_ and the Ser_201_ sidechain methylene (Fig. 3K). The ergoline A-ring makes van der Waals contacts with the methylenes of Met_209_ and Ser_201_ but the A-ring nitro group substituent does not engage in direct interactions with Sec14 (Fig. 3K).

Compared to the ergoline fused ring system, the tricyclic ABC himbacine rings occupy a shallower position in the lipid-binding pocket (compare Figs. 3B and C). The ABC-ring core takes advantage of an amphiphilic environment consisting of a hydrophobic surface formed by Tyr_122_, Tyr_151_, Leu_147_, Glu_150_, and a polar surface formed by Tyr_151_, Ser_173_, Thr_175_, Arg_208_, Met_209_ (Fig. 3F,I). The himbacine B- and C-rings are stabilized by hydrophobic interactions that primarily involve the C-ring and four Sec14 residues (Fig. 3F). The γ- butyrolactone A-ring is accommodated by the polar surface of the amphiphilic himbacine- binding pocket and its binding is stabilized entirely by polar interactions (Fig. 3I). In that regard, the proximity of the A-ring keto oxygen to Ser_173_ and Thr_175_ recapitulates the strategy by which the headgroup phosphate of the natural Sec14 ligand PtdCho is coordinated (Schaaf et al., 2008). The A-ring lactone oxygen is positioned in a polar environment provided by the Tyr_151_ hydroxyl of, and the backbone amides of Arg_208_ and Met_209_.

### Binding modes for distal SMI moieties

The distal SMI moieties are defined as: (i) NPPM/NPBB B-rings (Fig. 3J), (ii) the ergoline trichloroethyl-carboxylic ester group (Fig. 3K), and (iii) the himbacine linker and D-ring (Fig. 3L). All distal moieties adopt similar binding modes in the mostly hydrophobic subregion of the collective SMI binding envelope. Compared to the native ligand PtdCho, the distal moieties of NPPM/NPBB and himbacine overlap with the PtdCho glycerol backbone and proximal acyl chain-binding space. By contrast, the ergoline trichloroethyl-carboxylic ester group does not overlap PtdCho-binding space (Fig 2C).

With respect to specific interactions, the NPPM481 fluorobenzene B-ring is caged in a hydrophobic pocket formed by the sidechains of Thr_175_, Met_177_, Val_194_, Ala_197_, and Phe_212_ (Fig. 3J), and the distal pyrrolidine group of NPBB112 engages in van der Waals contacts with Met_177_, Val_194_ and Ala_197_ (Fig. S1A). The tricholoroethyl-carboxylic ester group of ergoline is accommodated in a hydrophobic region formed by Met_209_, Phe_212_ and Ile_240_. The latter two engage a chlorine atom in hydrophobic interactions with the Phe_212_ interface conforming to the parameters of a ‘face-on’ halogen-π interaction (Fig. 3K). Binding is further supported by polar interaction of the ergoline carbonyl oxygen with Ser_198_. The himbacine D-ring engages in hydrophobic interactions with Leu_147_, Met_177_, Ala_197_ and Tyr_193_. Polar character to the D-ring environment is provided by Glu_124_ and Tyr_193_ (Fig. 3L).

### Structural basis for Sec14 ability to accommodate different SMI chemotypes

Comparative analysis of the Sec14::SMI structures revealed the structural basis of Sec14’s ability to accommodate SMIs of different chemotypes. The lipid-binding pocket occupied by SMIs is spacious (∼3000 Å^3^; Sha et al., 1998). Extended linear arrangement of rings (NPPM/NPBB), compact fused ring structures (himbacine), and fused rings derivatized with ester-linked extensions (ergoline) are all accommodated within this cavity without a requirement for major Sec14 conformational adjustments. In the Sec14::NPPM244 complex, both the ligand itself and the interacting Sec14 residues superpose precisely onto NPPM481 and onto the cognate sidechains. Despite significant differences in the B and C-ring orientations of the Sec14-complexed NPBB12 and NPPM481, only Phe_212_ and Met_177_ exhibit noticeable rotameric shifts that enable van der Waals interactions with the NPBB112 B- and C-rings, respectively (Fig. S2A). The distinct chemotypes of NPPM/NPBBs and ergoline notwithstanding, there are no significant rearrangements of the corresponding binding motif sidechains in the NPPM vs ergoline bound states (Fig. S2B). The most pronounced sidechain rotameric change is observed in the Sec14::himbacine complex where the Tyr_151_ phenol ring is rotated ∼90° from its position in the other four Sec14::SMI complexes to promote a stacking interaction with the himbacine C-ring (Fig. S2C). This is accompanied by minor readjustments of Glu_150_ and Leu_147_. The modest extent and nature of conformational changes that accompany binding of SMIs with different chemotypes reports an impressive versatility of the Sec14 lipid-binding cavity chemical environment. This versatility primarily rests with the sidechains of residues that line the cavity. The majority of the polar backbone atoms are sequestered in intra-protein hydrogen bonds (Fig. S3) and are typically not involved in SMI interactions.

### Atomistic MD simulations reveal the highly dynamic nature of the Sec14::NPPM481 complex

The Sec14::SMI crystal structures describe SMI binding modes in an open Sec14 conformer. However, transitions between closed and open conformers is an essential aspect of the Sec14 lipid-exchange cycle (Ryan et al., 2007; Schaaf et al., 2008). Are SMI poses adjusted within the Sec14 lipid-binding cavity during transition from the open to the closed conformer? To gain insight into the conformational plasticity of SMI-bound Sec14 and the behavior of bound ligand, we conducted 2 μs atomistic molecular dynamics (MD) simulations using Sec14::NPPM481 as subject. For the remainder of this study, NPPM481 serves as model SMI. The justification is that NPPM481 is a potent inhibitor and is the best characterized Sec14 SMI from in vitro, in vivo and structure/activity relationship (SAR) perspectives (Nile et al., 2014). These simulations required the development of a CHARMM-compatible forcefield for NPPM481 (Figs. S4, S5; see Materials and Methods).

The MD data revealed that the complex is highly dynamic. In two independent 1 μs trajectories, the protein backbone underwent large-scale conformational changes as evidenced by the sharp transitions in backbone RMSD at 487 ns (Fig. 4A) and 20 ns (Fig. S6A), respectively. Significant RMSD fluctuations of bound ligand indicate that NPPM481 is not fully immobilized upon binding to Sec14. The frequencies of these fluctuations decrease once the Sec14 conformational transition takes place (Fig. 4A, Fig. S6A).

**Figure 4.**
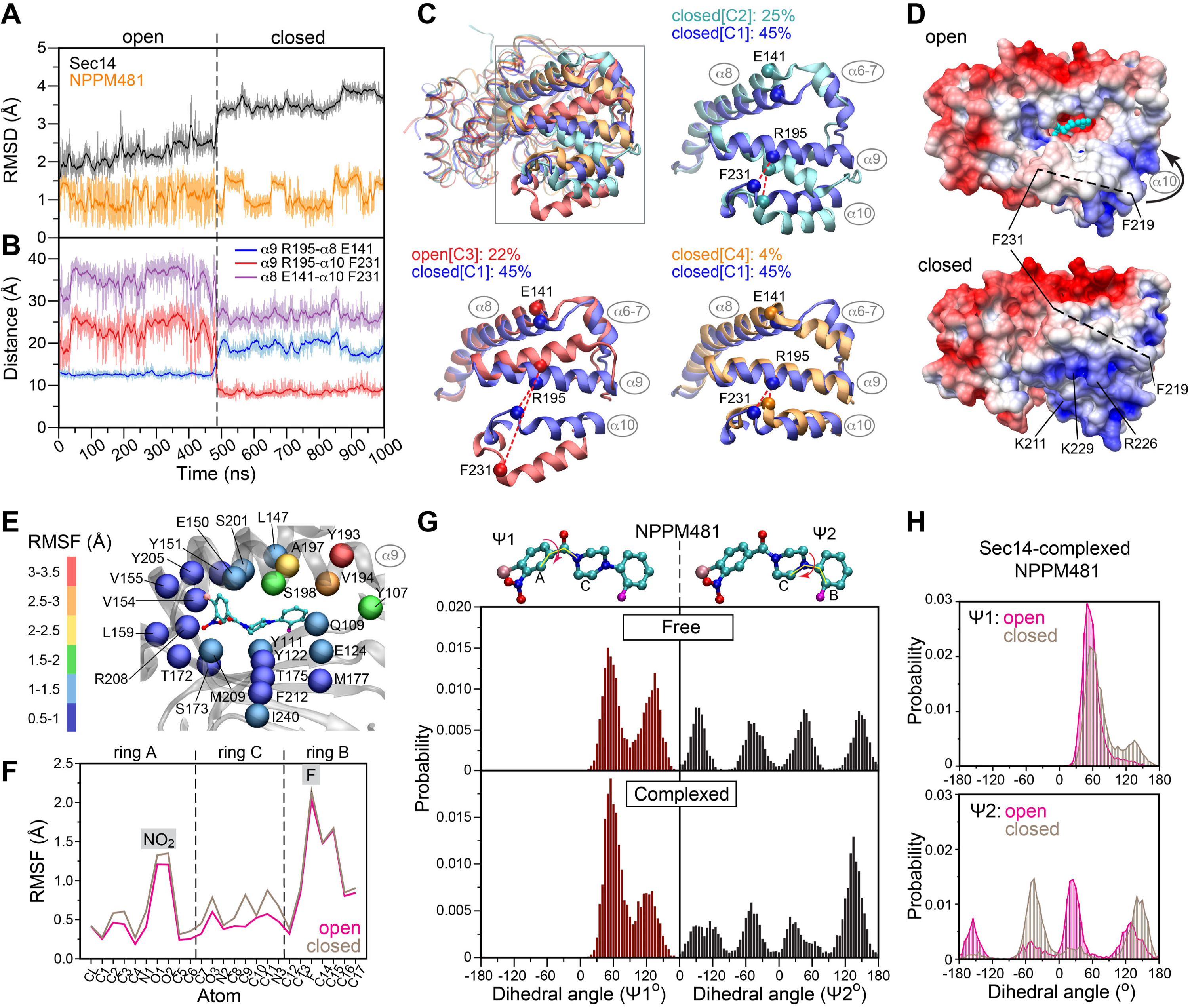
Dynamics of the Sec14::NPPM481 complex. **(A)** RMSD values of the Sec14 backbone (black) and NPPM481 (all heavy atoms, orange) plotted along the 1 μs trajectory. The open-to-closed transition at 487 ns is indicated with a vertical dashed line. **(B)** “Ruler” distances between the helices α8-α9 (blue), α9-α10 (red), and α8-α10 (purple). Open-to-closed transition of Sec14 involves the formation of the α9-α10 helix-helix interface. **(C)** Full and pairwise overlay of the representative Sec14::NPPM481 cluster structures illustrating structural variability of the helical segments α6-α10 The clusters are labeled C1 through C4. C1, C2, and C4 correspond to the closed conformation, while C3 corresponds to the open conformation. **(D)** Electrostatic potential mapped onto the representative cluster structures of the open and closed Sec14::NPPM481. The axis of the 219-231 α10 segment is shown with a dashed line. The realignment of the helical segments increases the hydrophilicity of the Sec14 surface through the formation of the positively charged patch over the ligand-binding site, and sequestration of the α10 hydrophobic face in helix-helix contacts with α9. **(E)** Backbone RMSF values of the Sec14 residues that line the lipid-binding pocket. The RMSF values are color-coded and mapped onto the corresponding Cα atoms. There is a gradient of flexibility along the lipid-binding pocket, with most dynamic residues residing on α9. **(F)** Per-atom RMSF values of the Sec14-bound NPPM481. The most dynamic segments are the nitro-group of the A-ring, and the entire B-ring that undergoes rotameric flips. The ligand dynamics is similar in the open and closed conformations of the complex. **(G)** Distributions of the dihedral angles Ψ1 (brown bars) and Ψ2 (black bars) that describe the relative orientations of rings A/C and C/B, respectively. The histograms were generated using 0.5 μs free NPPM-481 and 2 μs (combined) Sec14::NPPM481 production runs. Comparison of the Ψ1 and Ψ2 distributions between the free (top panel) and Sec14-bound NPPM481 (bottom panel) shows that the ligand, especially the B-ring, retains considerable rotameric flexibility upon Sec-14 binding. (H) Histograms of dihedral angles calculated using 1 μs trajectory of (A) and separated according to the open/closed states of the Sec14::NPPM481 complex. Transition to the closed state restricts the relative orientation of the C/B rings to two preferred values, Ψ2 ∼ -45° and ∼150°. The Ψ1 distribution is not significantly affected by the transition to the closed state.

Inspection of the MD trajectories revealed that the major contributor to the transition is repositioning of the Sec14 helical elements α9 and α10. Helix α10 comprises a “gate” element that controls lipid access to the binding pocket (Ryan et al., 2007; Schaaf et al., 2008). The “open” Sec14 conformation is represented in all crystal structures determined to date (Sha et al., 1998; Phillips et al., 1999), including the SMI complexes presented here, and is defined by a large spatial separation of helices α9 and α10. During the transition, α9 and α10 reposition to close the ligand-binding pocket. A convenient ruler to monitor this “open” to “closed” transition is the distance between the Cα carbons of Arg_195_ (α9) and Phe_231_ (α10) that reduces from ∼24 to ∼8 Å (Fig. 4B; Ryan et al., 2007; Schaaf et al., 2008). To provide a “static” reference point for helix motion, we chose the Cα carbon of Glu_141_ of the relatively immobile segment of helix α8. In the open conformation, α9 and α8 form an extensive helix-helix interface, while their respective distances to α10 fluctuate significantly due to the mobility of the latter (Fig. 4B, Fig. S6B). Upon transition to the closed state, α9 and α8 separate so that their inter-helical contacts are weakened while α9 and α10 reposition to form a new interface (Fig. 4B, Fig. S6B).

Cluster analyses of the combined trajectories provide a more detailed view of the conformational space sampled by Sec14::NPPM481 (Fig. 4C, Fig. S6C). Using a cutoff of 0.2 nm, we identify four clusters that cover 96% of the structures. Three correspond to closed Sec14 conformers (C1, 45%; C2, 25%; C4, 4%). Pairwise overlay of the representative cluster structures shows significant variability between them. These variabilities are highlighted by an “en bloc” movement of the α9/α10 helical pair in C2, and displacement of the α6-7 segment in C4. Cluster C3 (22%) corresponds to an open conformation with 24 Å separating helices α9 and α10 (Fig. 4C). We also note that Sec14 transition from open-to-closed conformers alters Sec14 surface properties. The corresponding surface electrostatic potentials show that Sec14 surface hydrophilicity increases upon α10 “gate” closure (as the α10 hydrophobic ridge is sequestered by formation of α9-α10 inter-helical contacts), and formation of a positively charged patch over the ligand-binding site (Fig. 4D).

Protein backbone RMSF analysis reveal a rich dynamic landscape for Sec14::NPPM481. In addition to the rearrangements of helices α6 - α10 identified by cluster analyses, the N-terminal segment of the G-module (a conformational switch element; Ryan et al., 2007), the 3_10_ helix T6, α5, and the loop that connects helices α4 and α5 (L_4/5_) show elevated RMSF values (Fig. S7A). The spatial fluctuations between these structural elements are apparent in the contact map that correlates the standard deviations of inter-residue distances (Fig. S7B). When viewed in the context of the 3D structure, the dynamic regions of Sec14 form a discrete structural element composed primarily of helical and loop regions (Fig. S7C).

Surprisingly, most of the residues that line the Sec14 ligand-binding cavity and directly interact with SMIs, show low RMSF values (<1.5 Å; Fig. 4E). Only five cavity residues score as highly dynamic (RMSF values >1.5 Å). Four of these lie in the C-terminal region of helix α9 (Tyr_193_, Val_194_, Ala_197_, Ser_198_). The fifth (Tyr_107_) marks the helix α5 terminus and the N-terminus of the G-module. Color-coding of the residues according to the RMSF values illustrates the dynamic “gradient” across the ligand-binding pocket (Fig. 4E). Bound NPPM481 mirrors this pattern. The A-ring, except for the -NO2 substituent, shows low RMSF values as it is tightly anchored to the Sec14 ligand-binding pocket (Fig. 4F). This is illustrated by the near-constant distances between the Cl substituent of ring A and the hydrophobic sidechains of Val_155_ and Tyr_151_ (Fig. S7D). By contrast, the NPPM481 B-ring is positioned near the mobile C-terminal segment of helix α9, and B-ring atoms show high RMSF values (Fig. 4F). Closure of the ligand-binding site brings α9 in proximity to the B-ring -- as reported by the reduction in distances between the apex of the B-ring, C-16, and the sidechains of Ala_197_ and Tyr_193_ (Fig. S7D).

To gain further insight into ligand dynamics, the distributions of two dihedral angles that report the relative orientations of the A-ring/C-ring (ψ1) and C-ring/B-ring (ψ2) were compared for free- and Sec-14 bound NPPM481 (Fig. 4G). While the ranges of sampled ψ1 and ψ2 values were similar between the two NPPM481 states, the Sec14-bound form showed clear preferences for ψ1 ∼ 60° and ψ2 ∼ 140°. Surprisingly, the B-ring rotameric flips manifested in the broad distribution of ψ2 values were not strongly attenuated in the Sec14::NPPM481 complex.

When the ψ1 and ψ2 distributions were independently analyzed for the open and closed conformers of the Sec14::NPPM481 complex, the data report that conformational transition to the closed state restricted the range of ψ2 by ∼100° and shifted the most frequently sampled ψ2 state from 25° to -45° (Fig. 4H). Significant restriction of NPPM481 rotameric states imposed upon closure of the ligand-binding site is also evident in RMSD traces where the ligand adopted a more limited set of discrete states in the closed conformer relative to the open conformer (Fig.4A, Fig.S6A). These data indicate NPPM481, particularly the B-ring, retains considerable rotameric flexibility in the Sec14::NPPM481 complex.

### NPPM481 displaces PtdCho from Sec14 in the absence of membranes

Sec14-targeted SMIs exhibit *C_log_P* values in a two-phase octanol/water system (*C_log_P* = log_10_([SMI]_octanol_/[SMI]_water_)) that predict their efficient partitioning into membrane bilayers. The NPPM481 *C_log_P* value of 2.7 estimates that, in an aqueous membrane bilayer system, ∼500 NPPM481 molecules are membrane incorporated for every molecule free in solution. Thus, it is suggested that SMIs intoxicate Sec14 by incorporating into the Sec14 lipid-binding cavity from a membrane environment during the lipid-exchange cycle (Nile et al., 2014). The current view is that Sec14 (and PITPs in general) shield the lipid-binding cavity from solvent when in solution, and that this cavity is largely inaccessible to solvated small molecules under those conditions (Smirnova et al, 2006). To test these hypotheses, we applied high-resolution ^19^F NMR spectroscopy to systems comprising PITPs, fluorinated SMIs, and isotropically tumbling lipid bicelles that provide the bilayer context and support PITP lipid transfer activity. A distinct advantage of NPPM481 for these studies is that it contains a fluorine atom that enables direct detection of the Sec14 binding process using ^19^F NMR.

First, a series of ^19^F NMR-detected binding experiments were conducted on an Sec14::PtdCho and NPPM481 binary system to determine whether NPPM481 can displace native ligand from the Sec14 binding pocket in the absence of membranes (Fig. 5A). To prepare the sample, recombinant phosphatidylglycerol (PtdGro)-bound Sec14 purified from *E.coli*, was populated with 16:0-18:1 PtdCho and re-purified to generate a monomeric Sec14:: PtdCho complex. ^19^F-PtdCho was subsequently exchanged into Sec14::PtdCho to ∼80% occupancy (see Materials and Methods). ^19^F NMR spectra were collected for two spectral regions. One was centered at -216 ppm to monitor ^19^F-PtdCho (Fig. 5B), and the second was centered at -123 ppm to monitor NPPM481 (Fig. 5C). Addition of progressively increasing amounts of NPPM481 to Sec14::^19^F-PtdCho results in drastic changes in both NMR spectra. Specifically, the ^19^F-PtdCho peak at -216.4 ppm decreases in intensity upon titration with NPPM481 -- indicating that ^19^F-PtdCho is being displaced from its position within the Sec14 pocket (Fig. 5B). This is accompanied by enhancement of the broad ^19^F spectral features in the -217.5 to -219.5 ppm range that we assign to the displaced ^19^F- PtdCho. Since there is no membrane environment present, the lipid likely remains associated with hydrophobic regions of the host Sec14 molecule. That ^19^F-PtdCho is indeed displaced by NPPM481 from its resident binding site in Sec14 is further corroborated by the increase in the intensity of the ^19^F peak at -125.9 ppm that corresponds to the Sec14::NPPM481 complex (Fig. 5C). The ^19^F signal originating from the complex is easily distinguishable from that of the free NPPM481 (-124.5 ppm), due to the broader line-width of the former associated with the increase in rotational correlation time.

**Figure 5.**
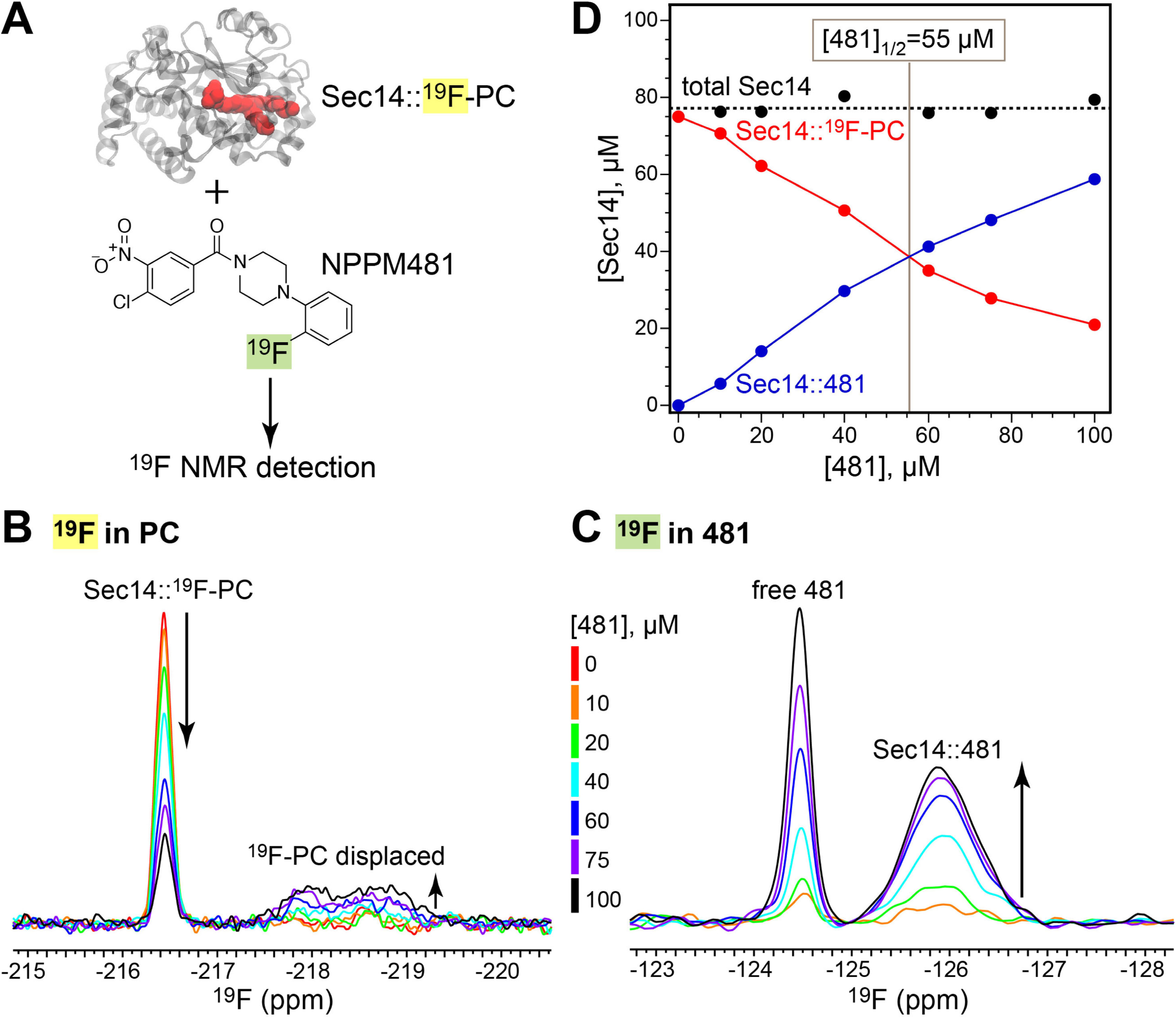
NPPM481 displaces PtdCho from Sec14 in the absence of membranes. PtdCho and NPPM481 labels are abbreviated to 481 and PC in this Figure and in Figs. 6-9 below. **(A)** Schematic representation of the ^19^F NMR-detected binding experiments in the binary Sec14:: ^19^F-PtdCho-NPPM481 system. The protein-bound ^19^F-PtdCho is shown in red. ^19^F NMR spectra showing the response of the ^19^F signals in ^19^F-PtdCho **(B)** and NPPM481 **(C)** upon varying the total NPPM481 concentration from 0 to 100 μM. The total Sec14::PtdCho concentration was 75 μM. **(D)** Redistribution of the PtdCho- (red) and NPPM481-complexed Sec14 (blue) upon increasing the total NPPM481 concentration. The data were obtained from the quantitative analysis of the ^19^F NMR spectra as described in Methods. Sec14 half-saturation is reached at the protein:NPPM481 ratio of 1:0.73 (i.e. 55 μM NPPM481).

Quantitative analysis of the ^19^F NMR spectra produced fractional populations of the Sec14 species complexed to ^19^F-PtdCho (f_Sec14::PC_) and NPPM481 (f_Sec14::481_). The plot of those values as a function of total NPPM481 concentration shows gradual redistribution between the ^19^F-PtdCho- and NPPM481-bound Sec14 species with increasing NPPM481 concentration (Fig. 5D). Moreover, the sum of the fractional populations is ∼1 for all points. This feature indicates that our quantitative analysis, conducted independently for the two spectral regions (see Materials and Methods), is internally self-consistent. Half-saturation of Sec14 with NPPM481 is achieved at a stoichiometry of ∼0.7 SMI:1.0 Sec14. Collectively, these results demonstrate a displacement of Sec14-bound ^19^F-PtdCho by NPPM481. Moreover, the data suggest that the Sec14 lipid-binding cavity is accessible to NPPM481 in an aqueous membrane-free system.

### Isotropically tumbling bicelles are suitable membrane mimics for investigating inhibition of Sec14 lipid-exchange activity

To establish the role of membranes in the displacement of PtdCho from Sec14 by NPPM481, we had to choose and validate a membrane mimic that is compatible with high-resolution NMR studies. Isotropically tumbling DiC14:0 PtdCho (DMPC):Di C6:0 PtdCho (DHPC) bicelles (q=0.5) are widely used in NMR studies of membrane proteins and presented an attractive option (Piai et al., 2017). To determine whether bicelles support Sec14-mediated lipid exchange, the fate of Sec14-bound ^19^F-PtdCho was monitored upon addition of bicelles. The ^19^F NMR spectrum of Sec14::^19^F-PtdCho prepared directly from purified PtdGro-bound Sec14 (see Materials and Methods) shows a peak at -216.4 ppm which corresponds to the ^19^F-PtdCho bound in the Sec14 lipid-binding cavity. The broad spectral features centered at ∼ -219 ppm correspond to the ^19^F-PtdCho peripherally associated with Sec14 (Fig. 6A). Upon addition of bicelles, the ^19^F spectrum changed dramatically -- resulting in the appearance of a single peak at -219.7 ppm. This peak can be unambiguously assigned to ^19^F-PtdCho transferred from Sec14 to bicelles based on the reference spectrum of ^19^F-PtdCho incorporated into bicelles during the preparation procedure (Fig. 6A, bottom panel). The transfer is also evident from the changes in the ^19^F-PtdCho dynamics quantified using ^19^F relaxation properties. ^19^F-PtdCho is more dynamic in bicelles than when bound to Sec14 as evidenced by the longer ^19^F longitudinal relaxation time of bicelle incorporated lipid (T_1_ =1.4 s) compared to Sec14-bound lipid (T_1_=0.8 s) (Fig. 6B).

**Figure 6.**
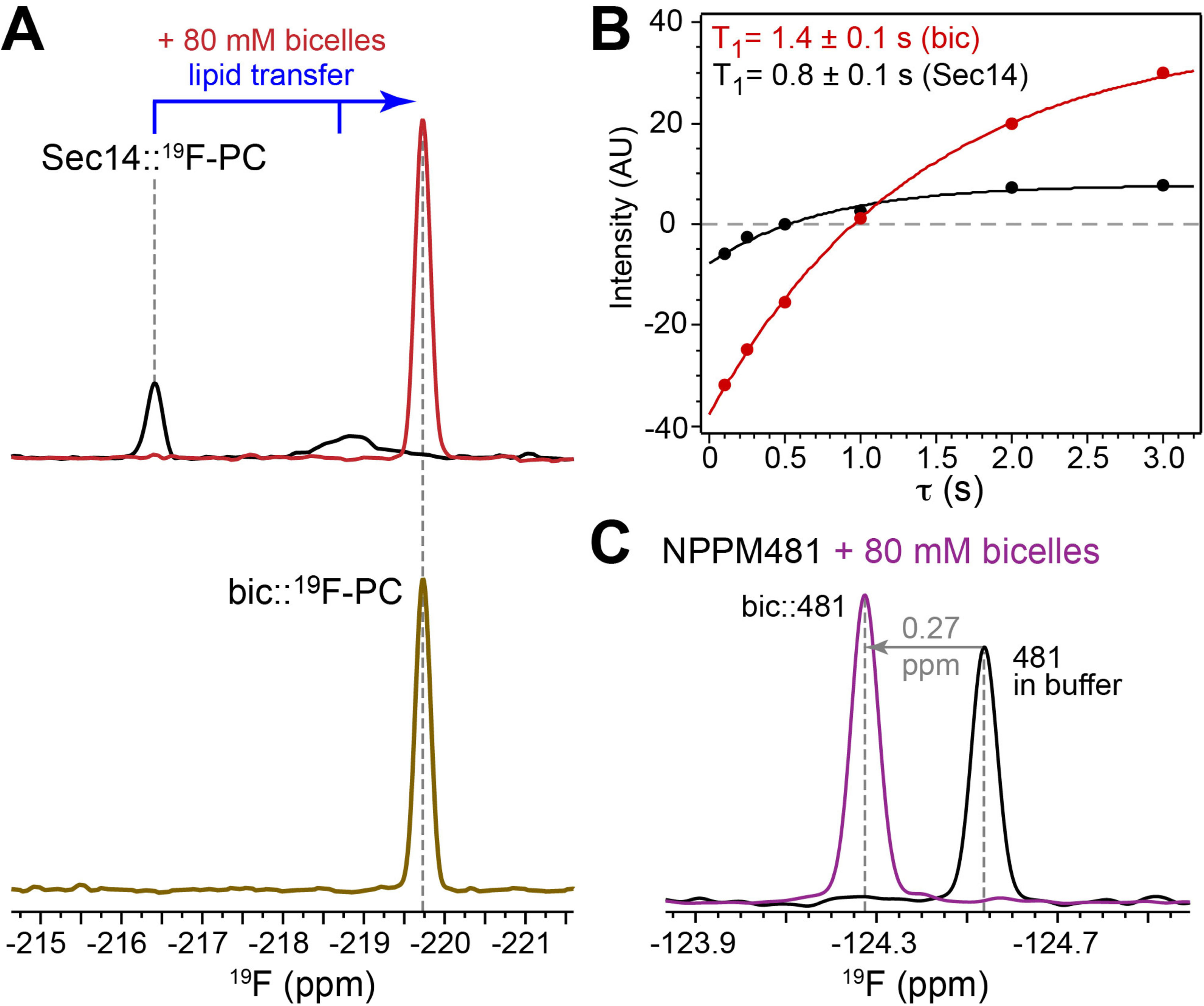
Bicelles support the lipid exchange function of Sec14 and provide a hydrophobic environment for NPPM481 partitioning. **(A)** Top panel: ^19^F NMR spectrum of the Sec14::^19^F- PtdCho complex (black) undergoes drastic changes upon addition of DMPC/DHPC bicelles (red). The appearance of new peak at -219.7 ppm indicates the transfer of ^19^F-PtdCho from Sec14 to bicelles. Bottom panel: the spectrum of ^19^F-PtdCho incorporated into bicelles during preparation is shown for reference (tan). **(B)** ^19^F inversion-recovery data and T_1_ values for the ^19^F-PtdCho bound to Sec14 (black) and bicelles (red). **(C)** ^19^F spectra of 75 µM NPPM481 in buffer (black) and in the presence of DMPC/DHPC bicelles (purple). The concentration of bicelles in all experiments was 80 mM (total lipid).

### Simulation of NPPM481 distribution in a bilayer system

To determine the interaction mode of NPPM481 with membranes, we recorded the ^19^F NMR spectrum of NPPM481 in the presence and absence of bicelles. Addition of bicelles results in a 0.27 ppm shift of the NPPM481 ^19^F peak -- strongly suggesting that NPPM481 is fully partitioning from the solution phase into the hydrophobic bicelle environment (Fig X2C, peak labeled bic::481). As Sec14 is a peripheral membrane protein, we assessed the insertion depth and orientation of NPPM481 in the membrane as an estimate of its accessibility to Sec14. To that end, atomistic MD simulations of binary systems comprising NPPM481 and Di14:0 PtdCho (DMPC) bilayers were conducted. DMPC was employed as bulk lipid in the simulations to match our experimental conditions.

To eliminate bias, four systems with different initial conditions were prepared. In system 1, one NPPM481 molecule was placed in solution 20 Å above the bilayer. In systems 2, 3 and 4, sixteen NPPM481 molecules were pre-inserted into the membrane at the bilayer center (system 2), the hydrocarbon region (system 3, eight molecules per leaflet), and the headgroup region (system 4, eight molecules per leaflet). All production runs were 500 ns. In system 1, the NPPM481 molecule spontaneously entered the bilayer at 47 ns and remained there for the entire duration of the production run (Fig. 7A). In systems 2-4, the membrane- initialized NPPM481 molecules redistributed within the bilayer and equilibrated within ∼100 ns (Fig. S8A). Thus, the last 400 ns of all trajectories were used for subsequent analyses.

**Figure 7.**
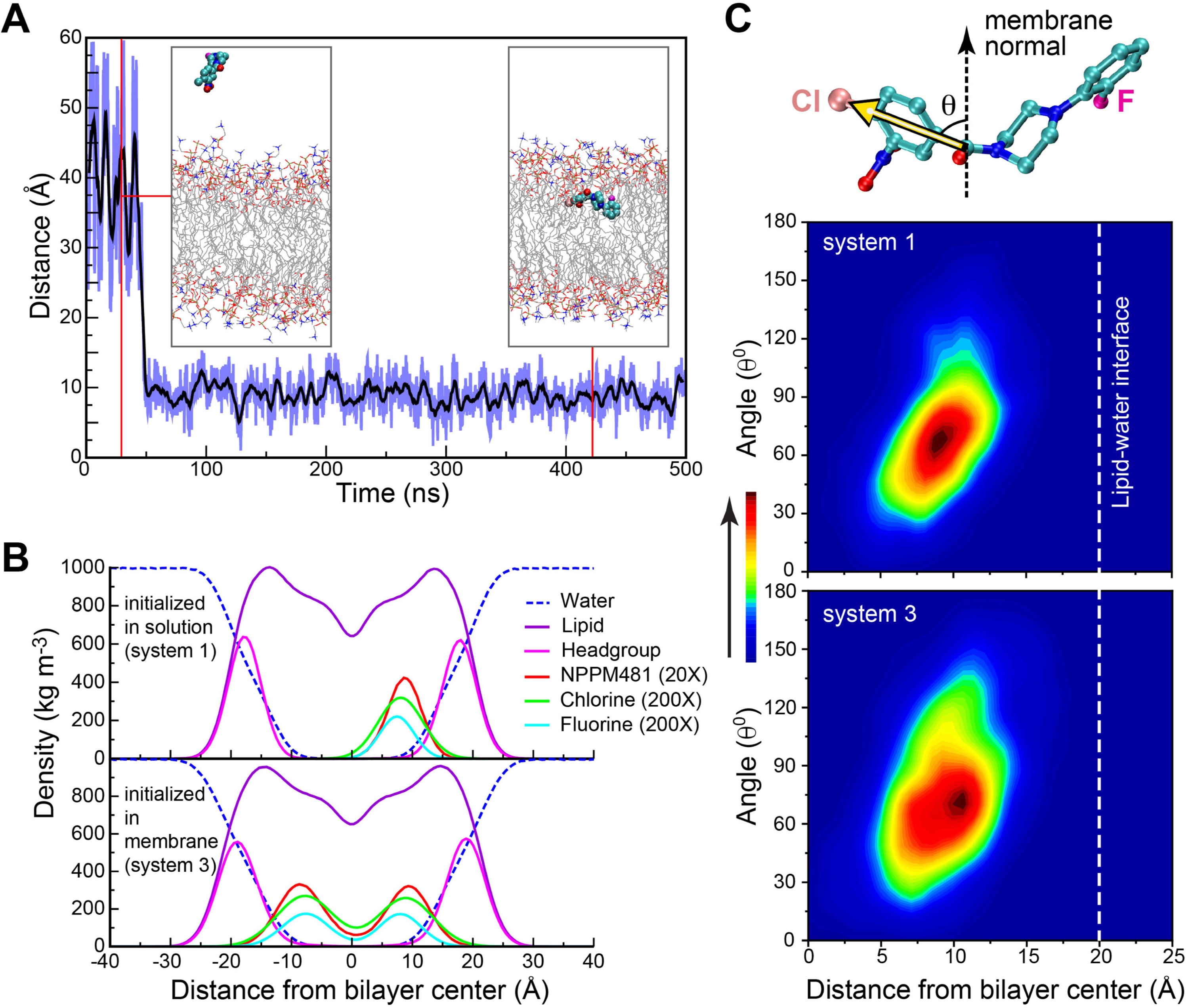
NPPM481 spontaneously partitions into membranes and positions itself below the head group region. **(A)** Distance along the Z axis (parallel to the membrane normal) between the NPPM481 center-of-mass and center of the membrane plotted along the 0.5 μs trajectory. The system was initialized with the NPPM481 molecule placed in solution 20 Å above the bilayer. Spontaneous partitioning into the membrane took place at 47 ns and persisted throughout the production run. **(B)** Mass density profiles calculated for the NPPM481 initialized in solution (top graph) and in the hydrocarbon region of the membranes (bottom graph). Irrespective of the initial conditions, NPPM481 molecules and their Cl and F substituents converge to the same position when projected onto the membrane normal. **(C)** Heatmaps correlating the distance between the NPPM481 A-ring and the bilayer center, and the A-ring tilt angle. The tilt angle is defined as the angle between the vector connecting the carbonyl carbon and the Cl atom of NPPM481, and the membrane normal. Irrespective of the initial conditions, the A-ring center is positioned ∼11-12 Å below the lipid-water interface, and the tilt angle is centered at ∼76°.

Irrespective of the initial conditions, NPPM481 molecules converge to the same position in the membrane as evidenced by the similarities of all mass density profiles along the membrane normal (Fig. 7B and Fig. S8B). The molecules partition into the hydrocarbon region of the bilayer with NPPM481 positioned ∼9 Å above the bilayer center and ∼9 Å below the headgroup region. The membrane densities of the halogen atoms (A-ring Cl, B ring F) report slightly deeper insertion with the corresponding peaks centered at ∼8 Å above the bilayer center in both cases.

To assess the orientation preferences of the NPPM481 A-ring in the membrane, we defined a vector that lies in the plane of the A-ring and connects the carbonyl carbon and the Cl atom of NPPM481. Angle θ between this vector and the membrane normal describes the orientation of the A-ring in the membrane (Fig. 7C). Representative heat maps for Systems 1 and 3, where θ is plotted against the distance between the A ring geometric center and the bilayer center, reveal a consistent pattern (Fig. 7C). Angle θ is centered at 76° (corresponding to a near-parallel A-ring vector orientation relative to the membrane surface), with a standard deviation of ∼30°. The distributions are modestly skewed in that smaller tilt angles correlate with deeper membrane insertion of the A ring, and vice versa. The A ring center is positioned at ∼9 Å away from the bilayer center and 11-12 Å below the lipid-water interface, with the closest approach to the interface of ∼6 Å. That is, a position where the ring center and its Cl substituent reside at the level of ester moieties of the headgroup region.

### Bicelles facilitate NPPM481-mediated PtdCho displacement from Sec14

The collective NMR and MD data report on the efficient partitioning of NPPM481 into the membrane bilayer. However, the MD data project an unexpectedly deep pose for NPPM481 in the bilayer environment. This raised the question of whether Sec14 is able to access membrane incorporated NPPM481. To determine how Sec14 interactions with NPPM481 are influenced by the presence of membranes, ^19^F NMR experiments were conducted using a ternary system comprised of Sec14::^19^F-PtdCho, NPPM481, and DMPC:DHPC bicelles (Fig. 8A). Bicelles were first added to the binary system containing 2-fold excess of NPPM481 relative to Sec14::^19^F-PtdCho with the respective concentrations set at 150 µM and 75 µM, respectively. Note that full displacement of ^19^F-PtdCho from Sec14 by NPPM481 was not achievable in the binary system. NPPM481 reaches its solubility limit at ∼120-150 µM and is no longer accessible to Sec14 at higher concentrations -- likely due to the formation of colloidal aggregates. Addition of bicelles produced distinct spectral changes (labeled 1 through 3) that report on the following three processes (Fig. 8B). First, the area of the ^19^F peak corresponding to the Sec14::NPPM481 complex increased by 25% -- indicating Sec14 is binding additional NPPM481. Second, the ^19^F peak of the unbound NPPM481 was shifted downfield by 0.27 ppm due to its quantitative partitioning into bicelles. We interpret the high intensity of the ^19^F bic::NPPM481 peak to reflect solubilization of NPPM481 aggregates (not detectable by NMR due to their high molecular mass) by bicelles. Third, both the ^19^F-PtdCho fraction remaining in the Sec14::^19^F-PtdCho complex and the fraction displaced by NPPM481, is transferred to bicelles. We conclude that NPPM481 partitions between two environments -- the Sec14 lipid-binding cavity and the bicelle, and that the SMI exhibits a higher apparent affinity for Sec14 than it does in the binary membrane-free system.

**Figure 8.**
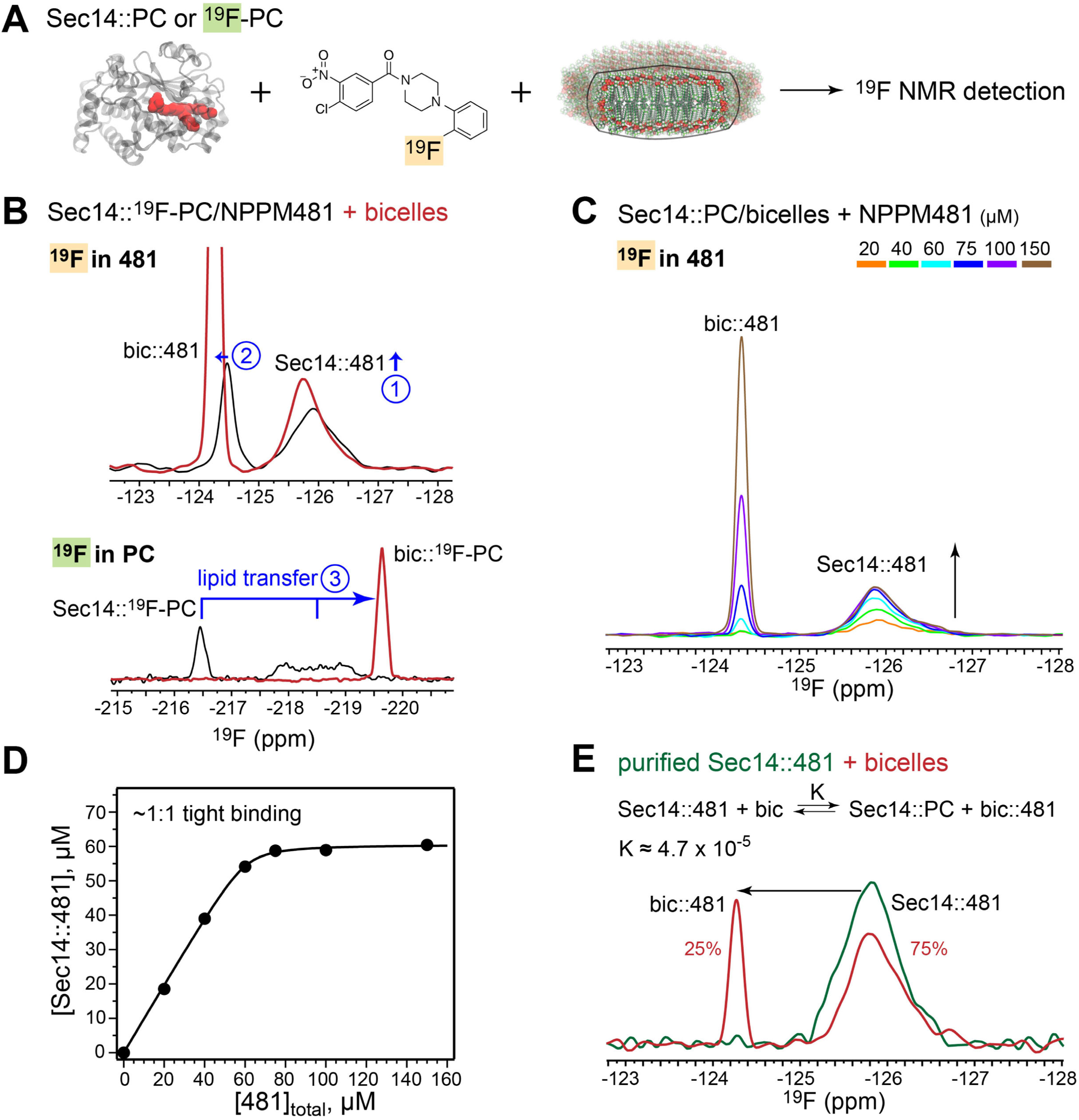
Membrane mimics facilitate the displacement of PtdCho by NPPM481 in Sec14. **(A)** Schematic representation of the ^19^F NMR-detected binding/displacement experiments in the ternary Sec14-NPPM481-bicelle system. The protein-bound PtdCho is shown in red. **(B)** Addition of bicelles to Sec14::^19^F-PtdCho and NPPM481 (molar ratio Sec14::^19^F- PtdCho:NPPM481=1:2) results in the 25% increase in the concentration of the Sec14::NPPM481 complex (1), partitioning of free NPPM481 into bicelles (2), and complete transfer of the ^19^F-PC from Sec14 to bicelles (3). **(C)** ^19^F NMR-monitored titration of Sec14::PC/bicelle system with NPPM481 (concentrations ranging from 0 to 100 μM). The buildup of the Sec14::NPPM481 complex is indicated with an arrow. **(D)** The lipid-NPPM481 displacement curve, plotted as a function of Sec14::NPPM481 concentration versus total NPPM481. The shape of the curve reports on high-affinity interactions of Sec14 with NPPM481, with full saturation attained under stoichiometric conditions. The solid line is shown to guide the eye. **(E)** Addition of bicelles to the purified Sec14::NPPM481 complex results in the substitution of PtdCho for NPPM481, with the resulting bic::NPPM481 and Sec14::NPPM481 populations of 25% and 75%, respectively. The estimated K value for the displacement reaction is 4.7×10^-5^. The total concentration of Sec14 is 45 μM.

To quantitatively estimate the ability of NPPM481 to displace ^19^F-PtdCho from Sec14 in the presence of membranes, we conducted ^19^F NMR-monitored NPPM481 titration experiments in a system where Sec14::^19^F-PtdCho was preincubated with 80 mM bicelles (Fig. 8C). The areas of the two ^19^F peaks, corresponding to bicelle-incorporated and Sec14- bound NPPM481 species, were used to calculate the concentration of the Sec14::NPPM481 complex and construct the lipid-NPPM481 displacement curve (Fig. 8D). Saturation is reached at ∼1:1 Sec14 to NPPM481 ratio -- reporting a high-affinity interaction. The tight binding regime precluded reliable determination of the equilibrium constant from these data. We therefore took advantage of the high-affinity interactions between Sec14 and NPPM481 to purify the Sec14::NPPM481 complex. Addition of bicelles to the complex resulted in only a modest exchange of bicelle-incorporated ^19^F-PtdCho for Sec14-bound NPPM481 with concomitant partitioning of the displaced NPPM481 from the Sec14 environment into bicelles (Fig. 8E). This was evident from the appearance of the bic::NPPM481 peak. By quantifying the ^19^F peak areas, we estimate the effective dissociation constant of the Sec14:: NPPM481 complex K_eff_ = K[bic]_t_ -- where K is the equilibrium constant for the exchange reaction and [bic]_t_ is the total concentration of lipids in bicelles (see Materials and Methods).

The K_eff_ and K values are 3.7 µM and 4.7·10^-5^, respectively. The low K value for the displacement reaction indicates that the formation of the Sec14::NPPM481 complex is thermodynamically strongly favored -- even when PtdCho is in an ∼2000-fold molar excess (80 mM total PtdCho versus 45 µM NPPM481; Fig. 8E). These data emphasize the high affinity of Sec14-NPPM481 interactions.

### Determinants of Sec14-NPPM481 high-affinity interactions

To probe high affinity Sec14-NPPM481 interactions in more detail, we took advantage of the close Sec14 paralog Sfh1 and two Sec14 mutants (Sec14^S173C^ and Sec14^V155F^). The common property is that all three proteins are resistant to inhibition by NPPM481.

Sec14^S173C^ is altered for a residue involved in both NPPM481 and PtdCho-headgroup binding – although PtdCho-binding/exchange activity is not strongly affected (Schaaf et al., 2008; Nile et al., 2014; Khan et al., 2016). Sec14^V155F^ alters a residue not involved in direct interaction with NPPM481. Rather, the Val → Phe substitution is identified in natural fungal polymorphisms of the Sec14 ‘VV-motif’ that render the PITP resistant to inhibition by NPPMs and ergoline (Khan et al., 2016: Pries et al., 2018). Sfh1 is naturally resistant to NPPM481 based on this polymorphism. Both the Ser_173_Cys and Val_155_Phe substitutions are presumed to directly inhibit SMI binding to Sec14 and Sfh1 (Khan et al., 2016; Khan et al., 2021). ^19^F NMR-based detection of the NPPM481-PtdCho exchange reactions enabled us to evaluate: (i) the significance of these specific substitutions in the formation of Sec14::NPPM481 complexes and (ii) the role of halogen atom substituent of the A ring in the NPPM SMIs.

To directly assess the ability of Ser_173_Cys and Val_155_Phe Sec14 mutants to extract membrane-partitioned NPPM481, we conducted the experiments in the “protein titration” mode. In that mode, increasing amounts of Sec14 proteins were added to a solution containing 75 µM of NPPM481 preincubated with 80 mM bicelles (Fig. 9A-C). Upon increasing protein concentration, the ^19^F peak of the bicelle-partitioned NPPM481 decreases in intensity due to the formation of the protein::NPPM481 complex. The latter is evident from the appearance of a broad ^19^F peak at ca. -(125.5-126.5) ppm. While 2-fold molar excess of Sec14 was sufficient to fully sequester membrane-partitioned NPPM481, the Ser_173_Cys and Val_155_Phe mutants failed to do so even when added at 4-fold molar excess.

**Figure 9.**
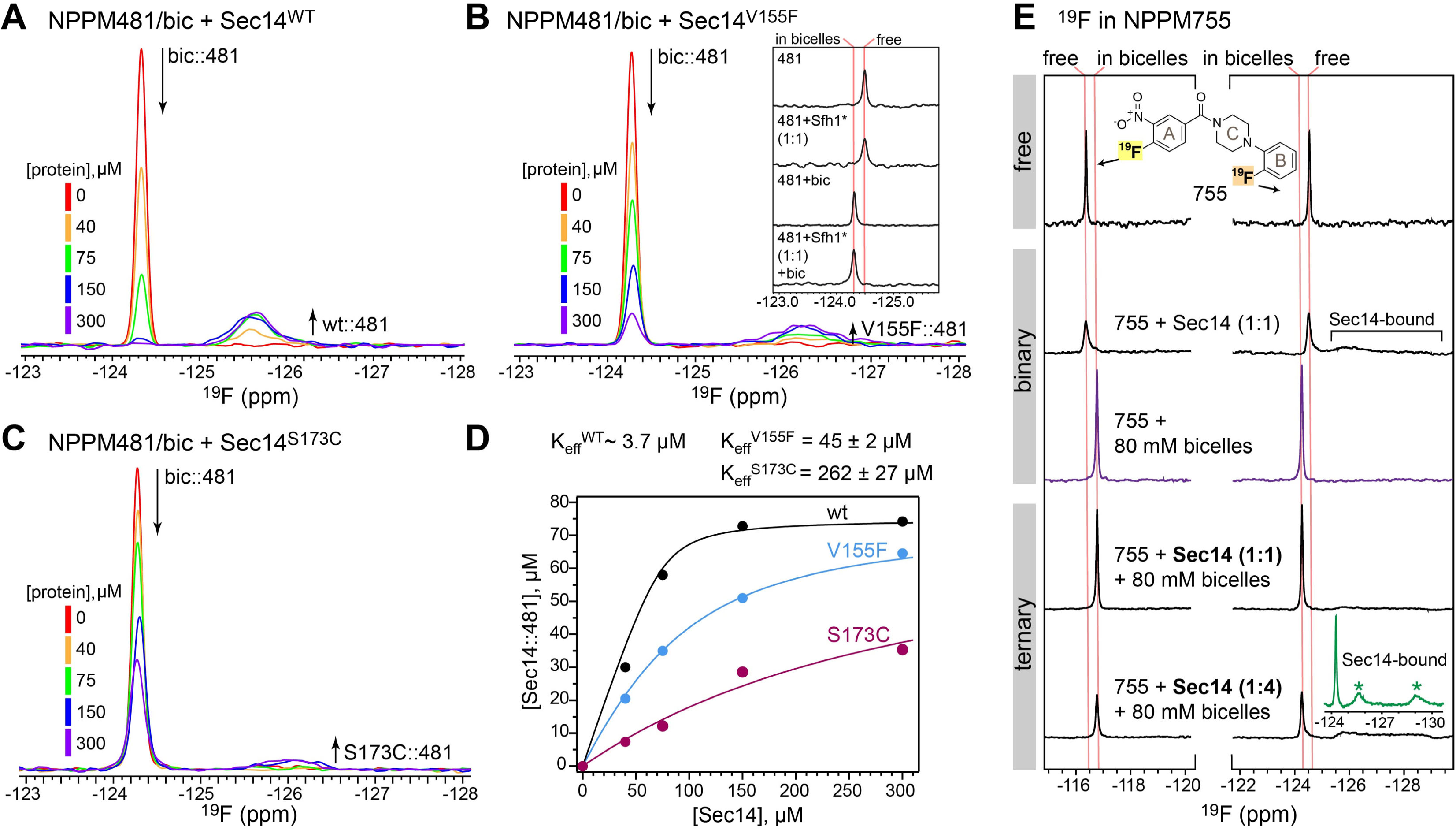
NPPM481 occupies the PtdCho headgroup-binding site. ^19^F NMR spectra of 75 µM NPPM481 in bicelles upon varying the total concentration of Sec14 variants: **(A)** wild- type, **(B)** Val_155_Phe, and **(C)** Ser_173_Cys, from 0 to 300 μM. The inset of (B) shows the ^19^F NMR spectra of 40 μM NPPM481 in solution, in presence of equimolar Sfh1*(Glu_126_Ala), and in presence of DMPC/DHPC bicelles. **(D)** The binding curves of NPPM481 to Sec14 wild type and variants (Ser_173_Cys and Val_155_Phe) in the presence of DMPC/DHPC bicelles. The K_eff_ values for the Sec14 variants were obtained by fitting the curves as described in Methods. For Sec14, the K_eff_ value was estimated using the experiment of Figure 8 (E). **(E)** ^19^F NMR spectra of 75 μM NPPM755 in solution, in presence of equimolar Sec14, in presence of DMPC/DHPC bicelles, in presence of equimolar Sec14 and bicelles, and 4-fold excess of Sec14 and bicelles. The chemical structure of NPPM755 is depicted in the inset of top panel showing locations of the probes. Inset of the last panel shows the spectrum collected with offset at -126.8 ppm with the spectral width of 10 ppm.

We used depletion of the bicelle-partitioned NPPM481 to construct the displacement curves and extracted the K_eff_ values (see Materials and Methods) to quantify the effects of these mutations on Sec14-NPPM481 interactions. Intrusion into the A-ring binding environment by the Val_155_Phe missense substitution resulted in a 12-fold decrease in the apparent affinity of NPPM481 to Sec14, as reported by the corresponding K_eff_ (V_155_F) value (Fig. 9D). For comparison, the exchange activated Sec14 paralog, Sfh1* that has a native Phe residue in the equivalent position (Schaaf et al., 2011), failed to bind NPPM481 either in the absence or presence of bicelles (inset of Fig. 9B). These data indicate that, while the residue identity at position 155 is an important determinant, it is not the sole determinant of the PITP affinity to the SMIs of this particular chemotype.

Ser_173_ is essential for maintaining the environment of the lipid-binding pocket and for ligand binding based on the engagement of its sidechain in: (i) polar interactions with the oxygens of the NPPM481 nitro group, and (ii) H-bond interactions with residues that also directly contact bound NPPM481 (Fig. S3). These structural data are consistent with demonstrations that a Ser → Cys substitution renders Sec14 insensitive to SMIs in vivo and in vitro (Nile et al., 2014; Pries et al., 2018). Indeed, the conservative nature of the Ser_173_Cys missense substitution notwithstanding, its effect on the Sec14-NPPM481 interaction was drastic -- resulting in a 70-fold decrease of apparent affinity (Fig. 9D).

To evaluate the role of the A-ring halogen substituent in the formation of the Sec14- SMI complexes, we investigated the interactions between NPPM755 and Sec14. NPPM755 belongs to the same chemotype as NPPM481, but has a fluorine substituent in the A ring instead of chlorine (Fig. 9E). The A- and B-ring fluorine atoms of NPPM755 give rise to distinct ^19^F signals at -116.4 and -124.6 ppm, respectively. In the binary system containing equimolar Sec14 and NPPM75, both ^19^F peaks broaden significantly due to the chemical exchange between unbound and Sec14-bound species. The latter manifests itself as the broad spectral features in the range of -(216-218) ppm. This behavior is qualitatively similar to that observed for the Sec14-NPPM481 binary system -- indicating that NPPM755 can also access the Sec14 lipid-binding pocket in the absence of membranes.

As expected, NPPM755 quantitatively partitions into bicelles in a binary NPPM755/bicelle system as reported by the chemical shift changes of both ^19^F peaks in those conditions (Fig. 9E, purple spectrum). The fractional population of the Sec14-bound NPPM755 species in a Sec14/bicelle/NPPM755 ternary system was estimated using the intensity decrease of the bicelle-partitioned NPPM755 ^19^F peak upon addition of Sec14 to a NPPM755/bicelle system. The fractional population of the Sec14::NPPM755 complex is 8% and 63% at ligand-to-protein ratios of 1:1 and 1:4, respectively (Fig. 9E). This estimates K_eff_ to be in the range of ∼200 μM. That is, a value similar to the K_eff_ of Sec14^S173C^-NPPM481 interactions (Fig. 9D). Despite the low affinity of NPPM755 binding, both ^19^F signals originating from the Sec14-bound NPPM755 could be detected with the appropriate adjustment (see Methods) of the NMR parameters (inset of Fig. 9E, green spectra).

## Discussion

Chemogenomic screens identify yeast Sec14 as a druggable protein (Hoon et al., 2008: Lee et al., 2014), and SMIs of multiple chemotypes have been validated to target Sec14 with exquisite specificity (Nile et al, 2014; Filipuzzi et al., 2016; Pries et al., 2018). Precisely how Sec14- directed SMIs engage and inhibit their target is not understood. Herein, we address unresolved questions regarding the mechanism of Sec14 inhibition by SMIs of four different chemotypes – NPPMs, NPBBs, ergolines, and himbacine. We report crystal structures of Sec14 complexed with each of these SMIs, and deploy a combination of molecular dynamics simulation and ^19^F- NMR spectroscopy approaches to describe/quantify key aspects of the Sec14, lipid and SMI dynamics that accompany Sec14 inhibition.

Taken together, the data: (i) offer a structural rationale for how these SMIs arrest Sec14-mediated lipid exchange, and (ii) outline a pathway for how SMIs engage Sec14 in high affinity interactions. Moreover, our studies reveal new insights regarding the uncharacterized Sec14 lipid-exchange cycle. These include projections that Sec14 inserts deeply into the cytosolic leaflet of a membrane during the lipid exchange process, and that the Sec14 substructures involved in gating the lipid binding pocket are more dynamic than previously believed when this PITP is disengaged from membrane surfaces and free in solution. The collective results not only demonstrate the utility of Sec14-directed SMIs for arresting the Sec14 lipid-exchange cycle at discrete stages, but also provide critical structural information to guide rational design of next-generation anti-mycotics directed against Sec14 PITPs of virulent fungi.

### SMIs compete directly with native ligand for binding to Sec14

The Sec14::SMI structures report that SMIs of all four chemotypes exhibit binding modes that exploit the amphiphilic substructure deep within the Sec14 lipid-binding cavity that hosts the natural ligand PtdCho, and that these compounds compete directly with PtdCho for binding to that site. The steric incompatibility of SMI-binding with accommodation of the PtdCho headgroup, the glycerol backbone and the proximal acyl chain-binding constituents of PtdCho fully account for why the SMIs inhibit the PtdCho-binding/exchange activity of Sec14. As the space occupied by PtdIns in the Sec14 lipid-binding cavity overlaps with that occupied by PtdCho in the glycerol backbone and acyl chain binding regions (Schaaf et al., 2008), SMI- binding is also sterically incompatible with PtdIns binding.

As demonstrated by Nile et al (2014), NPPM481 challenge results in an operationally irreversible inhibition of Sec14 activity in vivo (where PtdIns is an abundant cellular phospholipid) and strongly inhibits PtdIns exchange activity in vitro. We posit that the mechanism of inhibition of the PtdIns-transfer activity has a similar structural basis as that of PtdCho. Comparative structural analyses of Sec14::NPPM481 and lipid complexes of its paralog (Sfh1::PtdCho and Sfh1::PtdIns) support this view. The NPPM481 B- and C-rings invade the binding space that is occupied by the glycerol backbone/proximal regions of the PtdCho *sn*-2 acyl chain (Fig. S9A), or the proximal half of the PtdIns *sn*-2 acyl chain (Fig. S9B). By comparison, the ergoline fused ring structure mostly occupies the PtdCho glycerol backbone/phosphoester group binding space (Fig. S9C) and the PtdIns glycerol backbone/proximal region of the *sn*-2 acyl chain binding space (Fig. S9D). A common pattern among NPPM/NPBB/ergoline SMIs is the engagement of residues: Ser_201_, Tyr_205_, Arg_208_, and Met_209_ that belong to the C-terminal region of helix α9 and the adjacent loop region. These residues are not involved in the interactions with PtdCho, but are involved in the interactions with PtdIns in the Sfh1::PtdIns complex (PDB entry 3B7Z; Schaaf et al., 2008).

### Comparative analysis of Sec14-SMI interactions

The distinct amphiphilicity patterns of the SMIs are efficiently accommodated by the common binding environment. H-bond interactions do not dominate SMI binding profiles as the contact mode of each SMI analyzed in this study involves at most one H-bond interaction with protein. This feature neatly recapitulates what is observed for PtdCho-binding where the Sfh1::PtdCho structure forecasts only one H-bond interaction between Sec14 and bound PtdCho (Schaaf et al., 2008). Hydrophobic sidechains dominate the SMI binding environment and create subpockets (e.g., Val_154_, Val_155_) that are opportunistically invaded by NPPMs, NPBB112, and ergoline. Polar hydroxyl-containing amino acids (Ser, Thr, Tyr) provide amphiphilic character to the Sec14 lipid-binding cavity. Tyrosines are especially notable in this regard due to the impressive range of interactions they form with SMIs. These include H-bonding, ring stacking, methyl-π, and ‘edge-on’ halide**-π** interactions. Ser_173_, Thr_175_ and Ser_201_ similarly engage in multiple modes of interaction with PtdCho and SMIs (polar, H-bond, hydrophobic). Thus, the PtdCho-binding subregion of the Sec14 lipid-binding cavity provides a spatial arrangement of hydrophobic and hydrophilic surfaces that offers chemical complementarity to diverse sets of small molecules.

Interestingly, the Sec14 dynamics that accompany the transitions between open and closed Sec14 conformers that drive PtdIns/PtdCho exchange have only a minor effect on the interaction pattern of NPPM481 with Sec14. We speculate that the formation of high-affinity interactions between Sec14 and NPPMs (and likely NPPBs), is facilitated by the conformational flexibility of SMIs that permits sampling of interactions and channels the pose to an optimal fit.

### Ser173 -- an unusual case of a non-conservative Ser to Cys substitution

Ser_173_ is of particular interest as it is not only directly involved in all Sec14::SMI binding modes, but it is critical for productive inhibition of Sec14 activity in each case. Indeed, we find a subtle Ser_173_Cys missense substitution results in essentially complete Sec14 resistance to the presently validated SMIs in vitro and/or in vivo. Our NMR binding analyses quantify the defect as an ∼70-fold reduction in affinity of Sec14^S173C^ for NPPM481. What is the basis for such dramatic effects? We conclude Sec14^S173C^ represents an uncommon case of a non- conservative Ser → Cys substitution. Perhaps the lower electronegativity of sulfur relative to oxygen diminishes the H-bonding potential of Cys relative to Ser. Alternatively, the larger atomic radius of sulfur might: (i) create steric obstacles for accommodating SMIs in the Sec14 lipid-binding cavity, or (ii) alter the local binding environment by virtue of differences in Cys sidechain geometries (i.e. bond lengths, Cβ-S-H angles) relative to those of Ser (He and Quiocho, 1991). The non-conservative nature of the Ser_173_Cys substitution illustrates just how exquisitely tuned the Sec14-SMI interactions are.

### Implications for the Sec14 lipid exchange cycle

The Sec14 PtdIns/PtdCho-exchange cycle is the engine that drives PtdIns presentation to the PtdIns 4-OH kinases and thereby promotes PtdIns-4-P signaling. Yet, the protein and lipid dynamics associated with this cycle remain poorly understood. Prevailing models envision the solution form of Sec14 to represent a closed conformer that transitions to the open conformer after initial binding to the membrane surface. The NPPM481 studies challenge this simple view. NPPM481 distributes into two pools – a minor aqueous pool and a predominant membrane-incorporated pool (Figure 10, step 1). Our demonstration that NPPM481 accesses the Sec14 lipid binding pocket in the absence of membranes (i.e. from the aqueous pool), and displaces bound PtdCho from its natural binding site, suggest Sec14 undergoes appreciable conformational dynamics in solution, and that there is access to the lipid binding cavity in membrane-free contexts. Thus, one pathway for Sec14 intoxication by NPPM481 is via invasion of the lipid-binding cavity directly from solution (Fig. 10, step 2).

**Figure 10.**
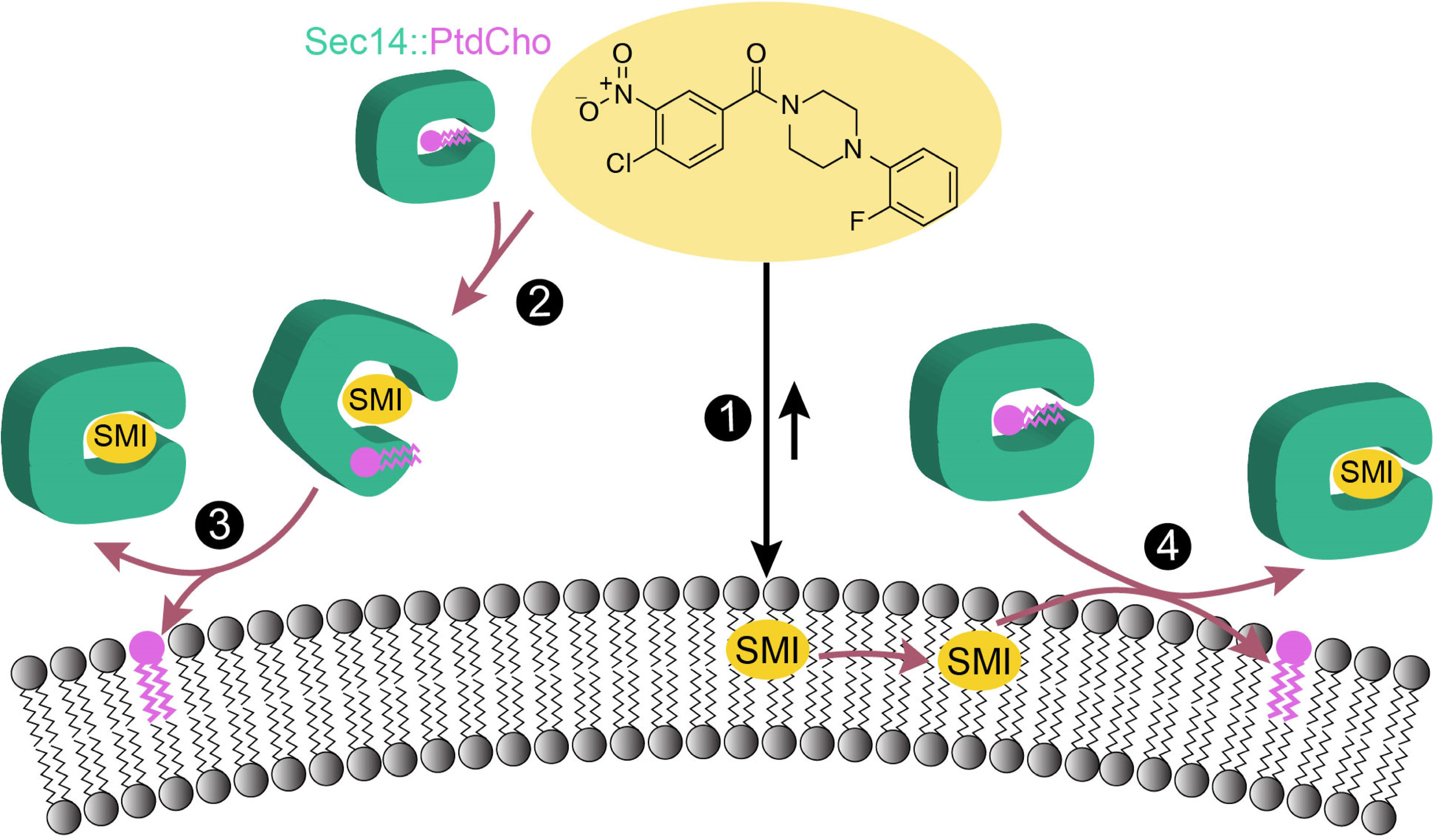
Pathways for Sec14 intoxication by small molecule inhibitors. NPPM481 is an amphipathic molecule that readily partitions into membranes from aqueous environments (1). NPPM481 binds Sec14 with high affinity and is able to invade the lipid binding cavity directly from solution and displace PtdCho from its natural binding site – resulting in formation of a soluble ternary complex (2). Upon engagement with membranes, the ternary complex resolves into an SMI-bound Sec14 complex with PtdCho released into the bilayer environment (3). The primary route of Sec14 intoxication is via a pathway where NPPM481 is incorporated into the Sec14 lipid binding cavity during a lipid exchange reaction (4).

The ternary Sec14/NPPM481/PtdCho complex formed in the absence of membranes is particularly intriguing as it fulfills the operational definition of an intermediate in the lipid exchange cycle. That is, upon addition of bicelles, the ternary complex releases the displaced PtdCho into the bilayer and cleanly resolves into an SMI-bound Sec14 product (Fig. 10, step 3). The ability of the ternary complex to complete a productive lipid exchange reaction upon bicelle addition raises the possibility that it is an open/partially open Sec14 conformer that initiates membrane binding -- rather than the closed Sec14 conformer that subsequently transitions to the open state to initiate lipid exchange after membrane binding. As a potential ‘trapped intermediate’ in the exchange cycle, the structure of this ‘arrested’ ternary complex now assumes significant interest. Our NPPM481 data highlight the utility of using SMIs as tool compounds to arrest, and to ultimately dissect, the exchange cycle in a stage-specific manner.

Although Sec14 can be intoxicated by SMI directly from solution, our data demonstrate NPPM481 exhibits a higher apparent affinity for Sec14 in bicelle-containing environments relative to the apparent affinity in binary membrane-free systems. We posit that membranes facilitate formation of the Sec481::NPPM481 complex by increasing the effective concentration of SMI by: (i) efficient solubilization and reduced dimensionality, and (ii) providing a thermodynamically favorable resolution to PtdCho displacement from its resident site in the Sec14 lipid-binding cavity. Thus, we conclude that the primary mechanism for Sec14 intoxication involves a reaction on the membrane surface where bound PtdIns/PtdCho is exchanged for membrane-incorporated SMI (Fig. 10, step 4).

In that regard, the MD experiments simulating NPPM481 partitioning into membranes project a deep pose for the SMI in the acyl chain region of the bilayer environment – one characterized by an orientation that is roughly parallel relative to the membrane surface. This configuration suggests that Sec14 must penetrate into the acyl chain region of the bilayer in order to access the SMI. As motion of the amphipathic α10 helix is a major contributor to Sec14 transitions between open and closed conformations, α10 is the most attractive candidate for insertion into the bilayer. Alternatively, Sec14 binding might locally disturb the bilayer such that the SMI rises closer to the membrane surface where it can then be accessed by Sec14.

### Insights into rational design of next-generation anti-mycotics

There is considerable urgency surrounding development of next-generation pharmaceuticals given the emergence of fungal ‘superbugs’ resistant to the limited set of currently available anti- mycotics for combating these infections (Perfect, 2017; Fisher et al., 2018). In that regard, the rigorous validation of Sec14 as specific target of SMIs of multiple chemotypes identifies fungal Sec14 orthologs as promising targets for development of new anti-fungal drugs. Whether a fungal Sec14 will display sensitivity to the presently validated SMIs is quite reliably predicted by two Sec14 elements – the PtdCho-binding signature (or barcode), and the Val_154_Val_155_ (VV)- motif that contributes to the contours of the PtdCho headgroup-binding environment (Khan et al., 2016; Khan et al., 2021; Bankaitis et al., 2022).

Interestingly, most virulent fungi express Sec14 PITPs with altered VV-motifs that substitute at least one of the Val residues with bulkier amino acids. Although such substitutions do not strongly affect the PtdIns/PtdCho-exchange activities of these proteins, our ^19^F NMR data demonstrate these alter the hydrophobic subpocket such that the capacity for NPPM/NPBB A-ring and ergoline carboxylic ester binding capacity is compromised. The fact that himbacine does not invade this subpocket accounts for why himbacine is the only validated SMI to date whose ability to inhibit Sec14 activity is insensitive to polymorphisms observed in the VV-motifs of Sec14 PITPs of virulent fungi (Danish Khan and V.A.B., unpublished data).

Thus, rational design strategies aimed at producing pan-fungal Sec14 SMIs could build on a himbacine fused ring scaffold as a strategy for circumventing VV-motif polymorphisms. That such strategies show promise is supported by the discovery of turbinmicin – a polyketide natural product that shows broad spectrum anti-fungal activity and is suggested to target Sec14 PITPs of pathogenic fungi that harbor polymorphisms in the VV-motif (Zhang et al., 2020).

The NPPM/NPBB A-ring-binding subpocket nevertheless remains an attractive element for exploitation in future design strategies as it is a rich site for mining SMI-Sec14 interactions. Such strategies would benefit from crystal structures of fungal Sec14::PtdCho complexes. As a case in point, our ^19^F NMR data indicate the close Sec14 paralog Sfh1 (a PtdIns/PtdCho exchange protein that harbors an altered VV-motif) fails to detectably bind NPPM481 -- even though incorporation of the Sfh1 Val_155_Phe substitution into the Sec14 context reduces apparent NPPM481-binding affinity only ∼12-fold. That the 12-fold reduction in apparent NPPM481 binding affinity is sufficient to endow Sec14^V155F^ with strong resistance to this SMI in vitro and in vivo reveals the problematically narrow dynamic range of assays currently used to monitor Sec14-SMI interactions. The power of ^19^F NMR spectroscopy in detecting weak PITP-SMI interactions recommends it as a facile tool for identifying/optimizing lead compounds for development of next-generation anti-mycotics.

## Materials and Methods

### Materials

1-palmitoyl-2-(16-fluoropalmitoyl)-sn-glycero-3-phosphocholine (^19^F-PtdCho), 1,2-dimyristoyl-sn-glycero-3-phosphocholine (DMPC), and 1,2-dihexanoyl-sn-glycero-3-phosphocholine (DHPC) were obtained from Avanti Polar Lipids (Birmingham, AL). α,α,α-trifluorotoluene was purchased from Sigma-Aldrich (St. Louis, MO). SMIs were purchased from ChemBridge Chemical Store (San Diego, CA.). The ergoline NGxO4 was kindly provided by Dominic Hoepfner (Novartis), and himbacine was obtained from Santa Cruz Biotechnology, Inc (Dallas, TX). All SMIs were dissolved in DMSO to final stock concentrations of 20-30 mM, and stored in the dark at −20°C.

### Crystallization of Sec14::SMI complexes

Octahistidine-tagged Sec14 (His8-Sec14) was purified from BL21-CodonPlus (DE3)-RIL cells (Stratagene, La Jolla, CA, USA) as described for Sfh1 (Schaaf et al., 2006) with minor modifications. Protein expression was induced with 60 µM isopropyl-β-D-thiogalactoside at 16°C for 20 h. Cells were lysed by glass bead beating in lysis buffer (300 mM NaCl, 25 mM sodium phosphate pH 7.5 and 5 mM β-mercaptoethanol), and the protein was affinity-purified from lysate using Ni-NTA affinity resin (Macherey-Nagel) and elution with imidazole. The His8-Sec14-enriched fractions were pooled and further resolved into two peaks by size exclusion chromatography (Superdex 75 16/600 column, GE Healthcare) at a flow rate of 1 mL/min in lysis buffer. Fractions of the slow eluting peak were pooled and concentrated to 5 mg/ml.

All crystallization experiments were conducted at room temperature using a sitting-drop vapor-diffusion method. The drop volumes were 2 µl and consisted of 1 µl Sec14::SMI or Sec14 solution as appropriate, and 1 µl of precipitant. Crystals appeared after 2-3 days of equilibration. To prepare Sec14::NPPM/NPBB complexes, 5 mg/mL His8-Sec14 was supplemented with 1 vol % of 20 mM SMI solution in DMSO. The Sec14::NPPM481 and Sec14::NPBB12 crystals were obtained after ∼2 days of equilibration under conditions where the precipitant was comprised of 129.5 mM sodium acetate, 64.8 mM Tris (pH 7.4), 4.6 % (w/v) PEG 4000, and 11.9 % (v/v) glycerol. The same procedures were applied to crystallize Sec14::NPPM244, except that the glycerol concentration was adjusted to 8.33 % (v/v) and pH of the precipitant solution was adjusted to 7.0 with acetic acid.

Crystals of the Sec14::ergoline complex were prepared in two stages. First, apo Sec14 was crystallized by diluting the His8-Sec14 stock to 2 mg/mL (25 µL of 5 mg/mL His8- Sec14, 30 µL 5x lysis buffer, and 5 µL lysis buffer). The precipitant composition was 170 mM sodium acetate, 85 mM Tris at pH 6.4, 25 % (w/v) PEG 4000, and 11.9 % (v/v) glycerol.

Second, an ergoline solution containing 0.45 µL precipitant, 0.45 µL 3.33 x lysis buffer, and 0.1 µL ergoline from a 30 mM stock in DMSO was prepared. A 1 µL aliquot of the ergoline solution was directly added to the 2 µL sitting drop containing apo Sec14 crystals in mother liquor followed by an overnight equilibration at room temperature.

To prepare crystals of the Sec14::himbacine complex, a 2 mg/mL His8-Sec14 solution prepared as described above for the first stage of the Sec14::ergoline crystallization was mixed with 1 vol % of 20 mM himbacine in DMSO. The crystals appeared after a ∼3-day equilibration against precipitant containing 170 mM potassium acetate, 85 mM Tris (pH 7.4, adjusted with acetic acid), 25 % (w/v) PEG 4000, and 3.6 % (v/v) glycerol.

### Structure determination

For data collection, Sec14::NPPM481, Sec14::NPPM112 and Sec14::NPPM244 crystals were transferred to a cryoprotecting solution (129.5 mM sodium/potassium acetate, 64.8 mM TRIS, 10 % (w/v) PEG 4000, 20 % (v/v) glycerol, pH 7.0) and flash frozen in liquid nitrogen. Sec14::ergoline and Sec14::himbacine crystals were directly flash frozen in liquid nitrogen. Diffraction data were collected at the 10SA (PXII) beamline at the Swiss Light Source (SLS) and indexed, integrated, and scaled using the XDS program package (Kabsch, 1993). The structures were solved using Molecular Replacement with PHASER (McCoy et al., 2007) using the Sec14::β-octylglucoside structure as search model (PDB entry 1AUA). Subsequent model building was performed in COOT and restrained refinement in PHENIX (Emsley and Cowtan, 2004; Adams et al., 2010). Ligand restraints were generated with Jligand (Lebedev et al., 2012). The quality of the final models was validated with the wwPDB Validation Server (https://validate-rcsb.wwpdb.org/). The coordinates of all structures were deposited in the Protein Data Bank. The accession numbers and statistics are given in Table S1 of the Supporting Information.

Structural analyses and figure preparation were carried out using Visual Molecular Dynamics (VMD) (Humphrey et al., 1996), UCSF Chimera (Pettersen et al., 2004), CCG Molecular Operating Environment (MOE) (Chem. Comp. Group Inc., Montreal, Canada), and LigPlot^+^(Laskowski & Swindells, 2011). Chemical structures of SMIs were generated in ChemBioDraw (PerkinElmer, Cambridge, MA).

### Computer simulations

#### NPPM481 parameterization with CHARMM force field

The initial set of CHARMM-compatible force field parameters was obtained with the CGENFF server (Vanommeslaeghe et al., 2010; Vanommeslaeghe and MacKerell, 2012) (Table S2), using the NPPM481 coordinates from the Sec14::NPPM481 crystal structure (PDB entry 7ZGC). Lone particle (LP) formalism (Pang et al., 2020) was used to describe the effect of the Cl ‘σ-hole’, a small positive charge associated with halogen bonds. All internal bond and angle parameters had a low penalty score (<1.5) and were taken as is. Partial charges with penalty scores >2.5 (Fig. S4A) and dihedral angles with penalty scores >10 (Fig. S5A) were selected for further optimization using the force field Toolkit (ffTK, version 2.1) (Mayne et al., 2013) distributed as the VMD plugin. NPPM481 parameterization followed the standard ffTK workflow and included optimization of geometry, partial charges, and dihedral angles. All quantum mechanical (QM) calculations were conducted using the DFT method (Hohenberg and Kohn, 1964; Kohn and Sham, 1965) implemented in Gaussian16 (Frisch et al., 2016). The following levels of theory/basis sets were used for the QM calculations: (i) B3LYP/6-311G++** -- initial geometry optimization, (ii) HF/6-31G* -- NPPM481-water interaction energies for charge optimization, and (iii) B3LYP/6-311G++** -- torsional scan profiles for dihedral angle optimization.

Partial charges were calculated from the NPPM481 water-interaction profiles (Fig. S4B,C), and optimized by fitting the molecular mechanics (MM) data to the QM-derived dipole moment, water interaction energies, and distances between water and each selected atom (Fig. S4D). Good agreement between the MM and QM target data is evidenced by low root mean square errors (r.m.s.e.) values for the water interaction energies and distances (Table S3). Optimization of dihedral angles relied on fitting the potential energy surfaces (PESs) generated by systematically scanning the dihedral angles and conducting QM calculations of energies for each NPPM481 conformation (Fig. S5B). For each of the four dihedral angles, quality of fit was evaluated based on the r.m.s.e. values and visual inspection. The optimized partial charges and dihedral angles of NPPM481 are given in Tables S4 and S5, respectively.

#### Molecular dynamics simulations of the Sec14::NPPM481 complex

All atomistic MD simulations were carried out using the Gromacs 2020.4 software package (Abraham et al., 2015) and CHARMM36 all-atom force field (Huang & MacKerell Jr, 2013). The crystal structure of the Sec14::NPPM481 (PDB entry 7ZGC) complex was used as the starting configuration for the MD simulations. The complex was solvated with TIP3P (Jorgensen & Jenson, 1998) water in a cubic box with an 8 nm edge. 43 Na^+^ and 35 Cl^-^ ions were added to create 100 mM salt concentration and neutralize the charge of the protein complex. After energy minimization, the system was equilibrated for: (i) 250 ps under an NVT ensemble at 310.15 K, (ii) 500 ps under an NPT ensemble with protein and ligand restrained; and (iii) 500 ps under an NPT ensemble with protein and ligand unrestrained.

Sec14::NPPM481 and solvent (water and ions) were coupled to separate temperature baths at 310.15 K using the V-rescale thermostat with a time constant of 0.1 ps. The pressure was controlled isotropically with the Berendsen barostat algorithm (Berendsen et al., 1984) during the NPT equilibration steps and the Parrinello-Rahman barostat algorithm during production runs (Parrinello & Rahman, 1981). Long-range electrostatic interactions were treated using the Particle Mesh Ewald (PME) method (Darden et al.,1993) with cubic interpolation and 0.16 nm grid spacing. Molecular bonds were constrained with the LINCS algorithm (Hess et al., 1997). The short-range neighbor list, van der Waals, and electrostatic cutoffs were all set to 1.2 nm. Periodic boundary conditions were used for all simulations. Randomized starting velocities were assigned from a Maxwell-Boltzmann distribution. Two independent trajectories of 1000 ns were simulated for the Sec14::481 system.

#### Molecular dynamics simulations of the NPPM481/membrane systems

Four NPPM481-membrane systems were assembled using the CHARMM-GUI membrane builder engine (Jo et al., 2008). In system 1, NPPM481 was placed 25 Å above the 64 x 64 DMPC bilayer surface. In systems 2-4, 16 NPPM481 molecules were placed at various positions in the 128 x 128 DMPC bilayer: bilayer center (system 2), hydrocarbon region (system 3, 8 molecules per leaflet), and the headgroup region (system 4, 8 molecules per leaflet). All systems were solvated with TIP3P water molecules in rectangular boxes having the dimensions of 6 x 6 x 12 nm (system 1) and 9 x 9 x 12 nm (system 2-4). 18 Na^+^/Cl^-^ (system 1) and 39 Na^+^/Cl^-^ ions (system 2-4) were added to create 100 mM salt concentration. Each system was energy-minimized and subjected to the multistage equilibration procedure as prescribed by CHARMM-GUI membrane builder engine. 500 ns production runs were executed for each system. Temperature and pressure were controlled using the Nose-Hoover (Hoover, 1985; Nosé, 1984) and Parrinello-Rahman semiisotropic coupling methods, respectively. All other algorithms were as described for the Sec14::NPPM481 simulations.

All MD data were analyzed using the following Gromacs analysis tools: *gmx rmsf* (RMSF analyses); *gmx mindst* (numbers of contacts between two selected atoms/groups), *gmx density* (membrane mass density profiles); *gmx distance* (distances between selected atoms/groups); *gmx gangle* (angles between the NPPM481 A-ring vector and bilayer normal); and *gmx cluster* (cluster analysis). Contact maps were generated using CONAN software package (Mercadante et al., 2018).

### Preparation of bicelle and protein samples for ^19^F NMR spectroscopy

All Sec14 NMR experiments were prepared in “NMR buffer 1” at pH 7.2 containing 100 mM NaCl, 25 mM Na_2_HP0_4_, 0.02% NaN_3_, and 8% D_2_O. The Sfh1* NMR experiments were prepared in “NMR buffer 2” at pH 7.2 containing 135 mM NaCl, 25 mM Na_2_HPO_4_, 0.02% NaN_3_, and 8% D_2_O.

#### Preparation of bicelles

Isotropically tumbling DMPC/DHPC bicelles (q=0.5) were prepared as previously described (Katti et al., 2020). In brief, the appropriate aliquots of DMPC and DHPC solutions in chloroform were dried under vacuum. For ^19^F-PtdCho containing bicelles, an aliquot of ^19^F- PtdCho solution in chloroform was added to create a molar ratio of 1:1067 ^19^F-PtdCho:(total DMPC+DHPC). The lipid films were resuspended by vortexing in “NMR buffer 1” to create a 1:2 DMPC:DHPC molar ratio and subjected to three freeze-thaw cycles to produce a clear and homogeneous bicelle solution. The stock solution prepared using this procedure contained 100 mM DMPC and 200 mM DHPC in case of pure DMPC/DHPC bicelles; and 75 µM ^19^F-PtdCho, 27 mM DMPC, and 53 mM DHPC in case of ^19^F-PtdCho containing bicelles. Total lipid concentration was quantified by phosphate assay (King, 1932).

#### Preparation of protein samples

Sec14, its variants Val155Phe and Ser173Cys, and the Glu126Ala variant of Sfh1 (Sfh1*) were expressed in *E. coli* BL21 (DE3) and purified as described in (Khan et al., 2016; Schaaf et al., 2008) with minor modifications. All protein constructs contained an 8xHistidine tag at the N-terminus. Sec14 and Sfh1 purified from *E. coli* are primarily bound to phosphatidylglycerol (PtdGro) with a minor fraction (∼10%) bound to phosphatidylethanolamine (PtdEtn), respectively. After the purification, Sec14::PtdGro and Sfh1::PtdEtn were either exchanged into their corresponding NMR buffers and used as is or, in the case of Sec14 subjected to additional steps, to prepare Sec14::PtdCho, Sec14::^19^F-PtdCho, and Sec14:NPPM481 complexes.

#### Preparation of PtdCho-bound Sec14

Purified recombinant Sec14::PtdGro (10 mg) was mixed with sonicated 17 mg POPC in 50 mL of buffer containing 300 mM NaCl, 25 mM Na_2_HPO_4_ at pH 7.2, and 1 mM NaN_3_. The mixture was incubated at 37°C for 45 min. Sec14::PtdCho was purified on the HisTrap HP column, followed by the gel filtration chromatography on the HiPrep^™^ 16/60 Sephacryl^®^ S-100 HR column. The purified Sec14::PtdCho was exchanged into the “NMR buffer 1” for further experiments.

#### Preparation of ^19^F-PtdCho bound Sec14

A 23 µL aliquot of 10 mg/mL ^19^F-PtdCho was dried under vacuum for 2 hours and resuspended in 600 µL of 75 µM Sec14::PtdCho solution in the NMR buffer 1 by gentle vortexing. The resulting mixture was incubated at room temperature overnight. We estimate that 80% of pre-bound PtdCho was replaced with ^19^F-PtdCho in Sec14, and will subsequently refer to this preparation as Sec14:: ^19^F-PtdCho. The estimate was obtained by comparing the integral ^19^F NMR peak area of ^19^F-PtdCho bound to Sec14 with that of the control NMR sample that contained a known amount of ^19^F-PtdCho in DMPC/DHPC bicelles.

#### Preparation of NPPM481-bound Sec14

A 100 µL aliquot of 0.5 mM Sec14::PtdGro solution in a buffer containing 300 mM NaCl, 25 mM Na_2_HPO_4_ at pH 7.2, and 1 mM NaN_3_ was mixed with 10 µL of 5 mM NPPM481 solution in DMSO and incubated for 5 minutes on ice. The mixture was resolved on a Superdex 75 10/300 gel filtration column with Sec14::NPPM481 eluting as a single peak. The peak fractions were pooled, exchanged into the “NMR buffer 1”, and concentrated to 45 µM for NMR measurements.

### 19F NMR experiments

All experiments were conducted on the Avance III Neo NMR instrument (Bruker BioSpin, Billerica, MA) operating at the ^1^H Larmor frequency of 600 MHz and equipped with a Prodigy cryoprobe. The temperature was calibrated using methanol-d4 and set at 25°C. 1D ^19^F NMR spectra were collected using the spectral width of 15 ppm and the ^19^F carrier frequency of either -123.0 or -216.0 ppm. The recycle delay was set to 3 s in all experiments. The number of scans was 2048 and 8192 for the Sec14::PtdCho-NPPM481 spectra of Fig. 5B and Fig. 5C, respectively; 4096 for the Sec14::PtdCho-NPPM481-bicelle spectra of Fig. 8C; and 1024 for wild-type and Sec14 variant spectra of Fig. 9A-C. All Sfh1* spectra shown in the inset of Fig. 9B were collected with 128 scans. The spectra of the NPPM755-containing samples (Fig. 9E) were acquired with: 128 scans for free NPPM755 in solution, 512 scans for NPPM755 in bicelles without Sec14, and 2048 scans for the remaining spectra. To resolve the Sec14-bound resonances of NPPM755 at 1:4 SMI:protein ratio (Fig.9E, inset), a spectrum was collected with 2048 scans, using spectral width of 10 ppm at ^19^F carrier frequency of -126.8 ppm. All data were zero-filled twice and Fourier-transformed using the 70 Hz Gaussian apodization function, except for the spectra shown in Fig. 5B and Fig. 5C, where the Gaussian line broadening of 30 and 100 Hz, respectively, were applied. Sfh1* (Fig.9B, inset) and NPPM755 (Fig. 9E) data were processed with the Gaussian line broadening of 60 and 20 Hz, respectively.

^19^F longitudinal relaxation time, T1, measurements were conducted using 1D inversion-recovery pulse sequence. The experiments were carried out on the Sec14-complexed and DMPC/DHPC bicelle-incorporated ^19^F-PtdCho, using the recycle delay of 7 s, relaxation delays ô of 0.1, 0.25, 0.5, 1.0, 2.0, and 3.0 s; and 1024 scans per spectrum. The data were zero-filled twice and Fourier-transformed using the 70 Hz Gaussian apodization function.

The T_1_ values were determined by fitting the data with the following equation:

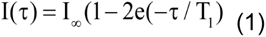

where I(ô) is the ^19^F peak area obtained at the relaxation delay ô, and Iϒ is the maximal peak area. The ^19^F chemical shifts were referenced to the external standard, α,α,α - trifluorotoluene (Sigma-Aldrich) that resonates at -63.72 ppm relative to CFCl3 at 0 ppm. All data processing and analysis was carried out using the MestReNova software package, v.14.2.0.

### NMR of binary Sec14::PtdCho-NPPM481 system

The interactions of Sec14-bound PtdCho with NPPM481 were monitored in a series of NMR-detected titration experiments, where increasing amounts of NPPM481 were added from the 7.5 mM stock solution in DMSO-d6 to the preparation of the Sec14:: ^19^F-PtdCho containing 75 µM total protein. The NPPM481 concentrations were 10, 20, 40, 60, 75, and 100 µM. The final sample dilution did not exceed 2%. The data were analyzed using the areas of ^19^F NMR peaks corresponding to the Sec14-bound ^19^F-PtdCho (-216.4 ppm); free NPPM481 (-124.5 ppm); and Sec14-bound NPPM481 (-125.9 ppm). At every NPPM481 concentration point, the fraction of Sec14 complexed to PtdCho, f_Sec14::PC_, was calculated from the ratio of the ^19^F- PtdCho NMR peak areas (PtdCho and NPPM481 are abbreviated to 481 and PC in all equations below):

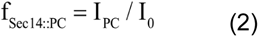

where I0 and IPC are the areas of the Sec14-bound ^19^F-PtdCho peak in the absence and presence of NPPM481, respectively. This calculation used the data of Fig. 5B. The fraction of NPPM481 bound to Sec14, f481,bound, was determined using the data of Fig. 5C:

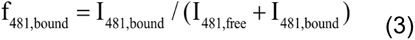

where I_481,free_ and I_481,bound_ are the areas of the ^19^F peak corresponding to the free and Sec14- bound NPPM481, respectively. The concentration of Sec14 complexed to NPPM481, [Sec14::481], was calculated as:

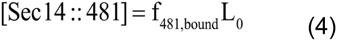

where [L]_0_ is the total concentrations of the NPPM481 in the sample. The fraction of Sec14 complexed to 481, f_Sec14::481_, was calculated as the ratio of [Sec14::481] to the total Sec14 concentration.

### NMR of ternary Sec14::PtdCho-NPPM481-bicelle system

The protein-NPPM481 binding experiments in the ternary system were conducted in the presence of the DMPC:DHPC bicelles (total lipid 80 mM), using two different types of experiments: (a) titration of Sec14::PtdCho/bicelles with NPPM481, and (b) titration of NPPM481/bicelles with Sec14::PtdGro and its two variants, Ser_173_Cys and Val_155_Phe.

In (a), the NPPM481 concentration was 20, 40, 60, 75, 100, and 150 µM. The concentration of the Sec14::481 complex was calculated using the spectra of Fig. 8C and equations Eqs.3 and 4, replacing I_481,free_ with I_bic::481_. The equilibrium constant for the exchange reaction:

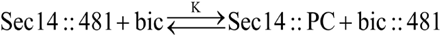

is defined as:

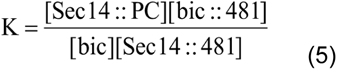

where [bic] and [bic::481] are the concentrations of bicelle lipids and bicelle-bound 481, respectively. Because the total lipid concentration in bicelles, [bic]t=80 mM is much larger than the total protein concentration, we can assume that [bic]Η[bic]_t_ and redefine the effective equilibrium constant as K_eff_=K[bic]_t_. The displacement curve, [Sec14::481] vs. L_0_, is described by the following equation:

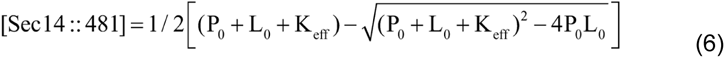

where P_0_ and L_0_ are the total concentrations of protein and NPPM481. The P_0_ was adjusted to 60 μM to account for active protein in the sample.

In (b), the concentrations of Sec14 variants were 40, 75, 150, and 300 µM, while the concentration of NPPM481 was kept constant at 75 µM. The concentration of Sec14::481 was calculated as:

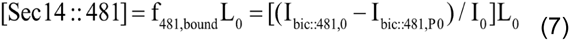

where L_0_ is a total concentration of NPPM481, I_0_ and I_bic::481_ are the ^19^F-NPPM481 peak areas in the absence and presence of protein, respectively. The binding affinity of NPPM481 to Sec14 variants Ser_173_Cys and Val_155_Phe was estimated by fitting the binding curve, [Sec14::481] vs. P_0_, using Eq. 6.

## Supporting information

Chen et al Supporting Information

## Data availability

The atomic coordinates and structure factors for all Sec14::SMI complexes are deposited in the PDB (https://www.rcsb.org) under the accession codes (SMI) as: 7ZGC (NPPM481), 7ZGD (NPPM244), 7ZGB (NPBB112), 7ZGA (NGxO4), and 7ZG9 (himbacine). Requests for further information should be addressed to the co-corresponding authors.

## Authors Contributions

The work and the experimental strategies were conceived by V.B., T.I.I., F.B. and G.S. The manuscript was written by T.I.I. and V.A.B with input from all authors. X.C. and S.S.K. performed the NMR experiments and analyzed the data with T.I.I. The NPPM481 forcefields were calculated by L.P., and L.P. and A.T. set up and executed the MD production runs.

Analyses of the MD data were performed by T.I.I., L.P., A.T., and V.A.B. A.H.N. identified and validated NPBB112 and himbacine as Sec14-directed SMIs and determined their IC_50_ values in PtdIns-transfer assays in vitro. Crystallization conditions were identified and optimized by P.J., and Z.H. collected and processed the X-ray data and built and refined the Sec14::SMI structures. Z.H., F.B., X.C., A.T., V.A.B, and T.I.I. analyzed the structural data.

S.S.K. devised methods to load Sec14 with ^19^F-PtdCho. S.M.G. optimized purification of the PtdCho-occupied Sec14 for these studies.

## Acknowledgements

This work was supported by grants NIH RO1 GM108998 and NIH R35 GM131804 to T.I.I. and V.A.B., respectively, and award BE-0017 from the Robert A. Welch Foundation to V.A.B. Z.H. and F.B. were supported by the European Research Council under the European Union’s Seventh Framework Programme (FP7/2007-2013), ERC grant agreement n° 310957 and the Deutsche Forschungsgemeinschaft (FOR2333). P.J. and G.S were supported by the Deutsche Forschungsgemeinschaft (DFG, German Research Foundation) under Germany’s Excellence Strategy – EXC 2070 – 390732324 and grant SCHA 1274/4-1 (to G.S.). We thank Dominic Hoepfner (Novartis) for providing the NGxO4 used in these studies, and acknowledge Benjamin Osborn (Biochemistry & Biophysics, Texas A&M University) for his initial involvement in optimizing methods to occupy recombinant Sec14 purified from *E. coli* with PtdCho. We also thank Jae Hyun Cho (Biochemistry & Biophysics, Texas A&M University) for helpful comments critical reading of the manuscript.

