## Supplementary material for "Mechanisms by Which Small Molecule Inhibitors Arrest Sec14 Phosphatidylinositol Transfer Protein Activity": Chen et al Supporting Information

**[2]Max Planck Institute for Developmental Biology**
**Spermannstrasse 35**
**72076 Tübingen, Germany**

**[3]Institute for Crop Science and Resource Conservation**
**Universität Bonn, Karlrobert-Kreiten-Strasse 13**
**53113 Bonn, Germany**

**[4]Department of Molecular & Cellular Medicine**
**Texas A&M University**
**College Station, Texas 77843 U.S.A.**

**Running Title:** Sec14 inhibition by small molecules
**Key Words:** Sec14 PITPs/ phosphoinositides/ protein and lipid dynamics/ lipid exchange/anti-mycotic drugs

**† -** Co-corresponding authors

### FIGURES

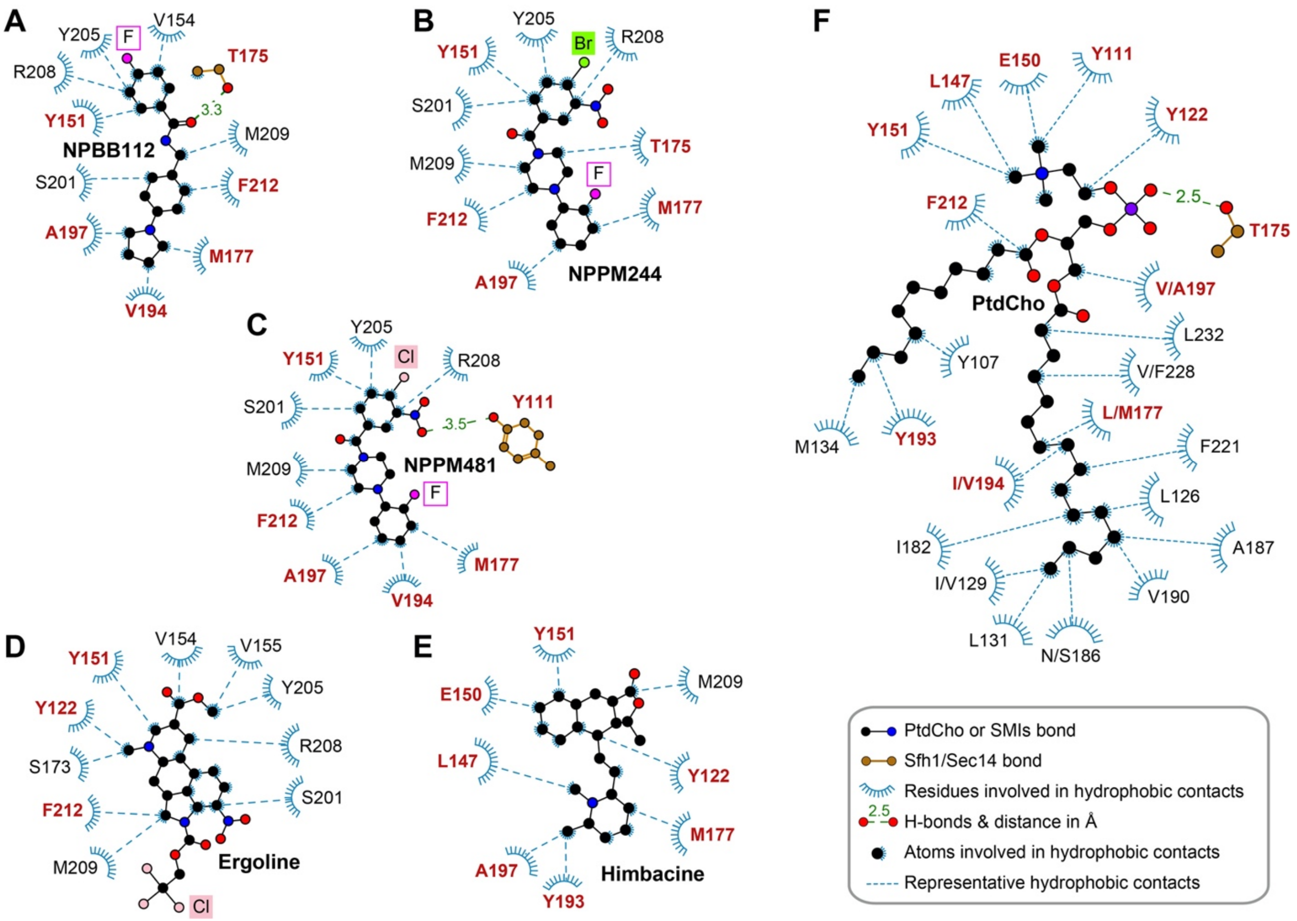


**Figure S1. SMI-specific interactions in the Sec14::SMI complexes.** LigPlot+ diagrams of the Sec14 complexes: **(A)** Sec14::NPBB112 (7ZGB), **(B)** Sec14::NPPM244 (7ZGD), **(C)** Sec14::NPPM481 (7ZGC), **(D)** Sec14::Ergoline (7ZGA), and **(E)** Sec14::himbacine (7ZG9). The cutoff for the hydrophobic interactions is set at 4 Å. Halogen atom interactions are not shown. **(F)** LigPlot+ diagram of the Sfh1::PtdCho complex. The residue numbering is adjusted to match the Sec14 numbers. There are three positions in the lipid-binding pocket where the amino acid identity between the two proteins differs: Leu[Sfh1]/Met[Sec14]$_{177}$, Ile[Sfh1]/Val[Sec14]$_{194}$, and Val[Sfh1]/Ala[Sec14]$_{197}$. Sec14 residues that are involved in interactions with both SMIs and PtdCho are highlighted in red font.

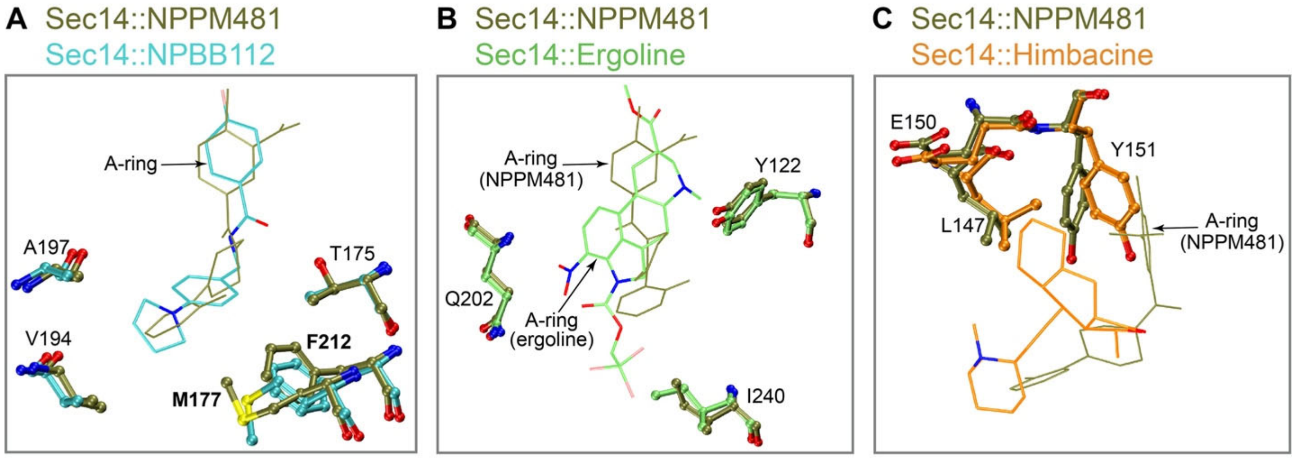


**Figure S2. Sec14 binding to the SMIs of different chemotypes is accompanied by small adjustments in the ligand-binding site.** Pairwise superposition of the Sec14 complexes: **(A)** Sec14::NPPM481 (tan) and Sec14::NPBB112 (cyan); **(B)** Sec14::NPPM481 (tan) and Sec14::ergoline (green); and **(C)** Sec14::NPPM481 (tan) and Sec14::himbacine (orange). Atom-specific coloring is applied to NPBB112, ergoline, and himbacine, but not to NPPM481. SMIs and selected Sec14 residues are shown in stick and ball-and-stick representations, respectively. In (A), while A-rings of NPPM481 and NPBB112 complexes superimpose well, the B-ring of NPBB112 is shifted ~90 degrees relative to that of NPPM481. $Phe_{212}$ and $Met_{177}$ sidechains rotate to accommodate that difference. All five Sec14 residues shown in (A) interact with B-rings of SMIs. In (B), $Tyr_{122}$ and $Ile_{240}$ of Sec14 interact with ergoline but not with NPPM481. $Gln_{202}$ contributes to the polar environment of the ergoline A ring. In (C), residue $Leu_{147}$ interacts with himbacine but not with NPPM481. $Tyr_{151}$ interacts with both himbacine and NPPM481, but its sidechain and those of $Leu_{147}$ and $Glu_{150}$ are rotated significantly relative to their poses in the Sec14::NPPM481 structure.

A

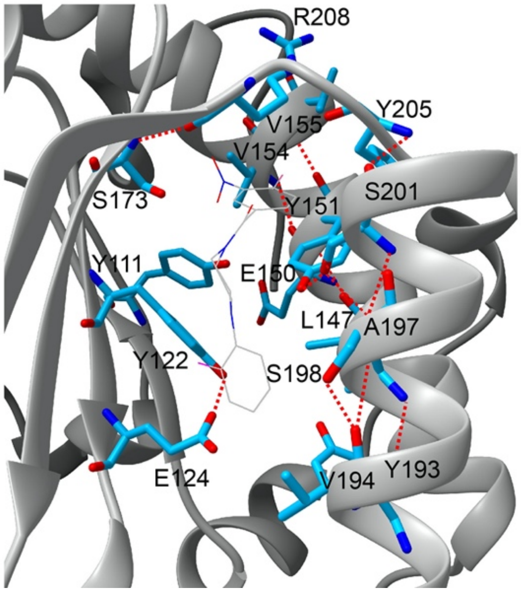


B

| | | | Sec14 complexes | | |
|---|---|---|---|---|---|
| | | Protein H-bonds | NPPM481 | Ergoline | Himbacine |
| Gm/β3 | 1 | Y111 (OH)-S173 (OG) | - | - | + |
| | 2 | Y122 (OH)-E124 (OE2) | + | + | + |
| | 3 | Y151 (NH)-L147 (O) | + | + | + |
| | 4 | V154 (NH)-E150 (O) | + | + | + |
| | 5 | V155 (NH)-Y151(O) | + | + | + |
| | 6 | S173 (NH)-R208 (O) | + | + | + |
| β3/Gm | 7 | S173 (OH)-R208 (O) | - | + | - |
| | 8 | A197 (NH)-Y193 (O) | + | + | + |
| | 9 | S198 (NH)-V194 (O) | + | + | + |
| | 10 | S198(OH)-V194 (O) | + | + | + |
| | 11 | S201 (NH)-A197 (O) | + | + | + |
| | 12 | S201 (OH)-A197 (O) | + | + | + |
| | 13 | S201 (OH)-Y151(Osc) | + | + | + |
| | 14 | Y205 (NH)-S201 (O) | + | + | + |
| β-turn | 15 | R208 (NH)-Y205 (O) | - | + | - |

**Figure S3. Sec14 residues that line the lipid-binding pocket form an extensive network of intra-protein H-bonds.** H-bonding analysis was carried out using Chimera software package. The criteria of distance and angle constraints were relaxed by 0.4 Å and 25 degrees, respectively. **(A)** Schematic representation of H-bonds in the Sec14::NPPM481 complex, with bound NPPM481 shown with thin lines. **(B)** List of all intra-protein H-bonds formed by residues involved in SMI interactions. 'Osc' in entry #13 denotes the hydroxyl oxygen of the Tyr sidechain. 'Gm' stands for the G-module element. The presence/absence of a given H-bond in a Sec14 complex is indicated with +/- symbols. The H-bonding patterns are very similar among the complexes.

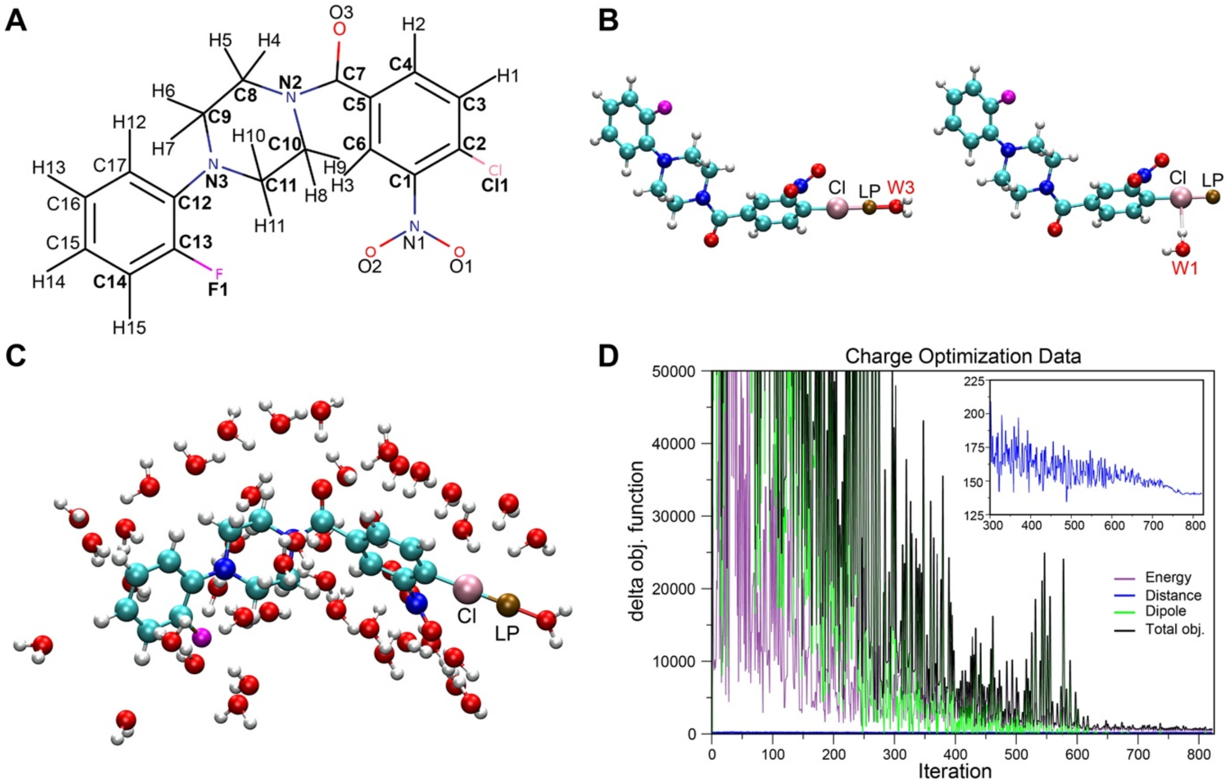


**Figure S4. Generation of CHARMM-compatible force field: optimization of partial atomic charges of NPPM481. (A)** Chemical structure of NPPM481 with atomic numbering. Atoms shown with boldface were subjected to charge optimization, as implemented in the ffTK plugin of VMD. **(B)** Two possible geometries of Cl-water interaction sites. Cl, the "lone pair" (LP), and water molecules are color-coded pink, brown, and red, respectively. **(C)** All water-interaction sites of NPPM481 used in the quantum mechanical calculations of water interaction energies and subsequent charge optimization. **(D)** The convergence of the total objective function and its individual contributions from the energy, distance, and dipole moment components during the charge optimization routine. The distance convergence plot is shown in the inset.

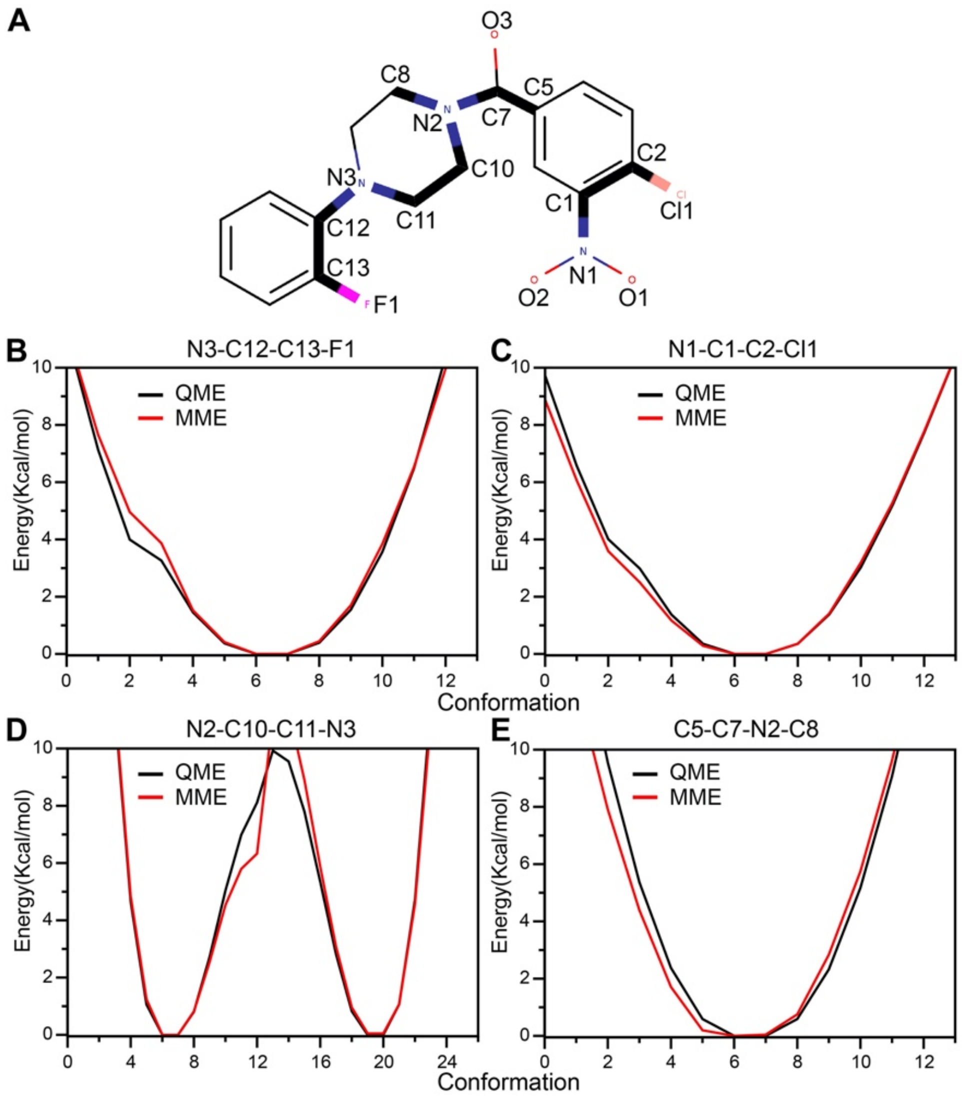


**Figure S5. Generation of CHARMM-compatible force field: optimization of NPPM481 dihedral angles. (A)** Four dihedral angles of NPPM481 that were subjected to optimization. Each angle is defined by four atoms whose connectivity is schematically shown with thick lines. **(B)-(E)** Results of the final refinement optimization for the four dihedral angles. The refined molecular mechanics (MM) parameters produce a good agreement between the MM (red) and the target QM (black) potential energy surfaces.

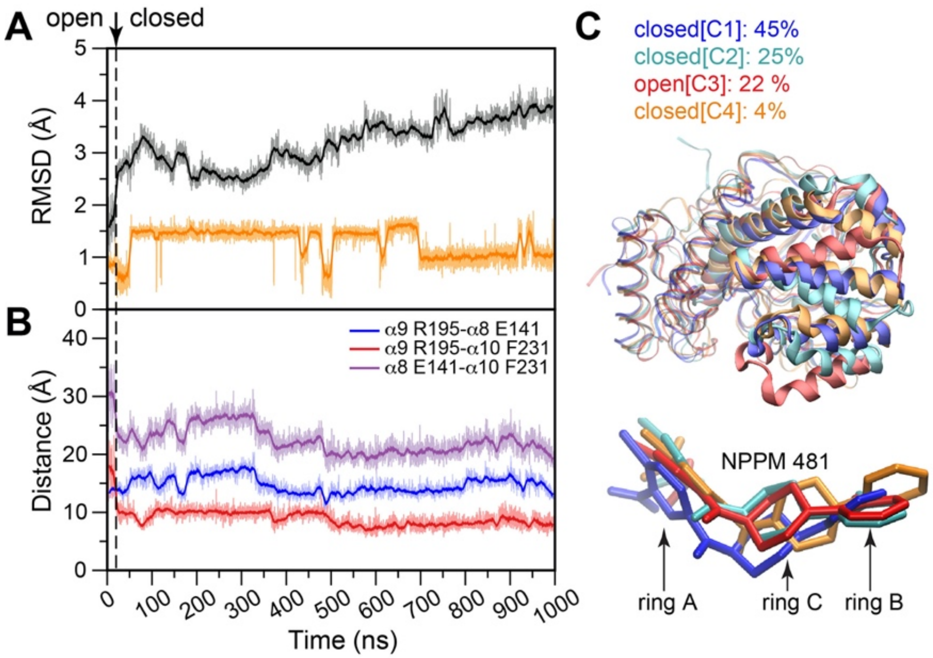


**Figure S6. Dynamics of the Sec14::NPPM481 complex. (A)** RMSD values of the Sec14 (backbone, black) and NPPM481 (all heavy atoms, orange) plotted along the 1 μs trajectory (replica 2). The open-to-closed transition occurs at 20 ns and is indicated with a vertical dashed line. **(B)** "Ruler" distances between the helices α8-α9 (blue), α9-α10 (red), and α8-α10 (purple). Open-to-closed transition of Sec14 involves the formation of the α9-α10 helix-helix interface. **(C)** Representative cluster structures of the Sec14::NPPM481 complex obtained from the analysis of the combined 2 μs production run. The overlay of Sec14-bound ligands, color coded according to cluster identity, is shown in the bottom panel. The four clusters, C1 through C4, cover 96% of the structures.

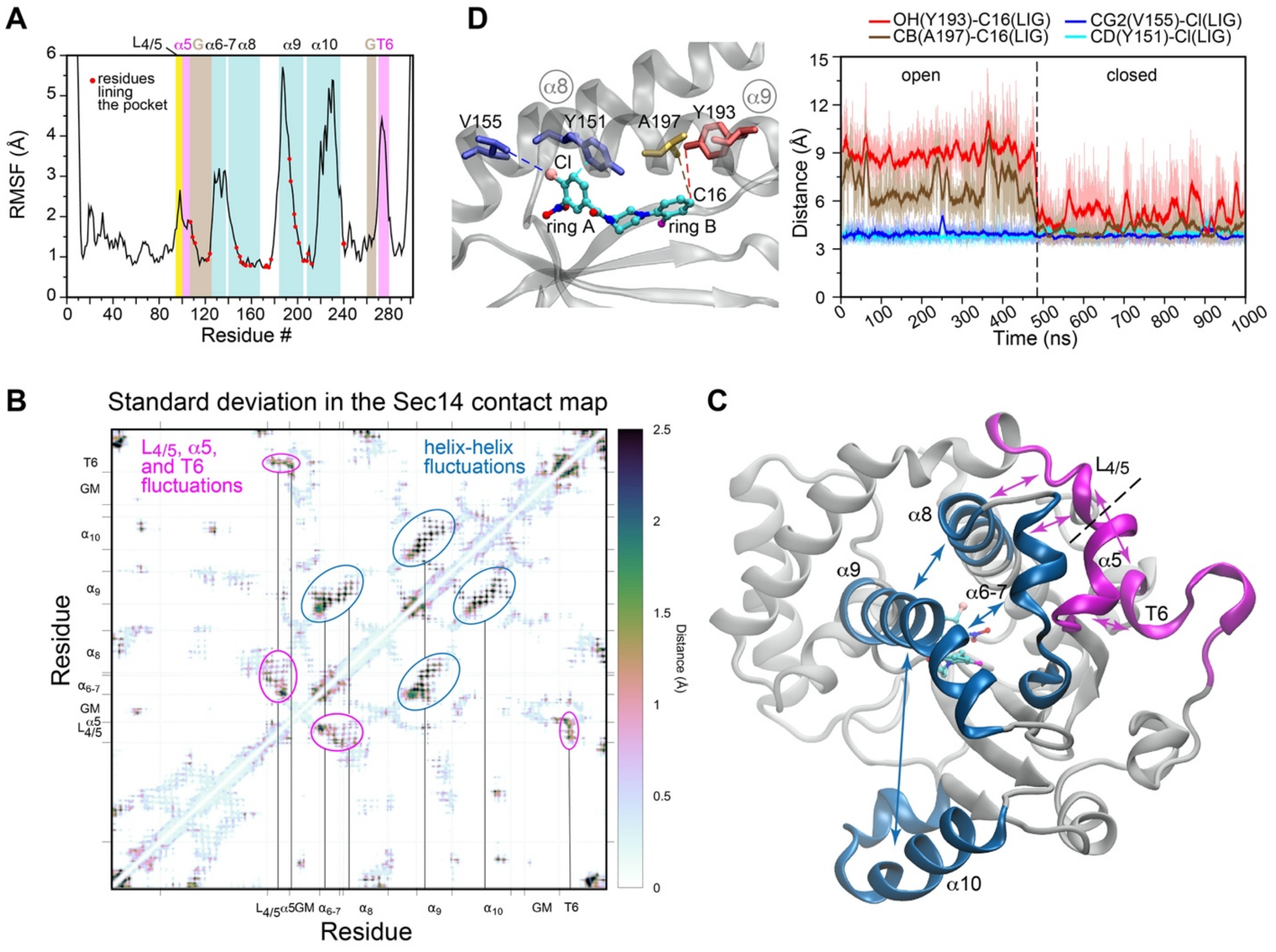


**Figure S7. Dynamics of the Sec14::NPPM481 complex. (A)** Sec14 backbone RMSF values calculated using the combined 2 μs trajectory. The structural elements with elevated RMSF values include the α-helical elements: α5, α6-7, α9, α10, and the N-terminal region of α8; the N-terminal segment of the G-module; the $3_{10}$ helix T6; and the loop that connects helices α4 and α5, $L_{4/5}$. Red circles mark the residues that line the lipid binding pocket. Of those, the most dynamic residues are in the N-terminal segment of the G-module and the C-terminal region of α9. **(B)** Fluctuations in the inter-residue contacts, expressed as RMSD values and calculated over the 1 μs trajectory (replica 1). The map was generated using CONAN software package, using the distance cutoff of 10 Å. The fluctuations in the helical contacts and α5/T6/$L_{4/5}$ are highlighted with blue and magenta circles, respectively. **(C)** Dynamic regions of Sec14 mapped onto the 3D structure of the Sec14::NPPM481 complex (7ZGC). Double-headed arrows schematically show the fluctuations observed in the CONAN plot. **(D)** NPPM481 is tightly anchored to Sec14 via the interactions of its A-ring. This is illustrated by the near-constant distances between the Cl atom of the A ring and the sidechain atoms of $Val_{155}$ and $Tyr_{151}$. The flexibility of the B ring and the α9 helix manifests itself in the fluctuating distances between the C16 atom of NPPM481 and the sidechain atoms of $Ala_{197}$ and $Tyr_{193}$. There is significant decrease of those distances upon the closure of the ligand-binding pocket.

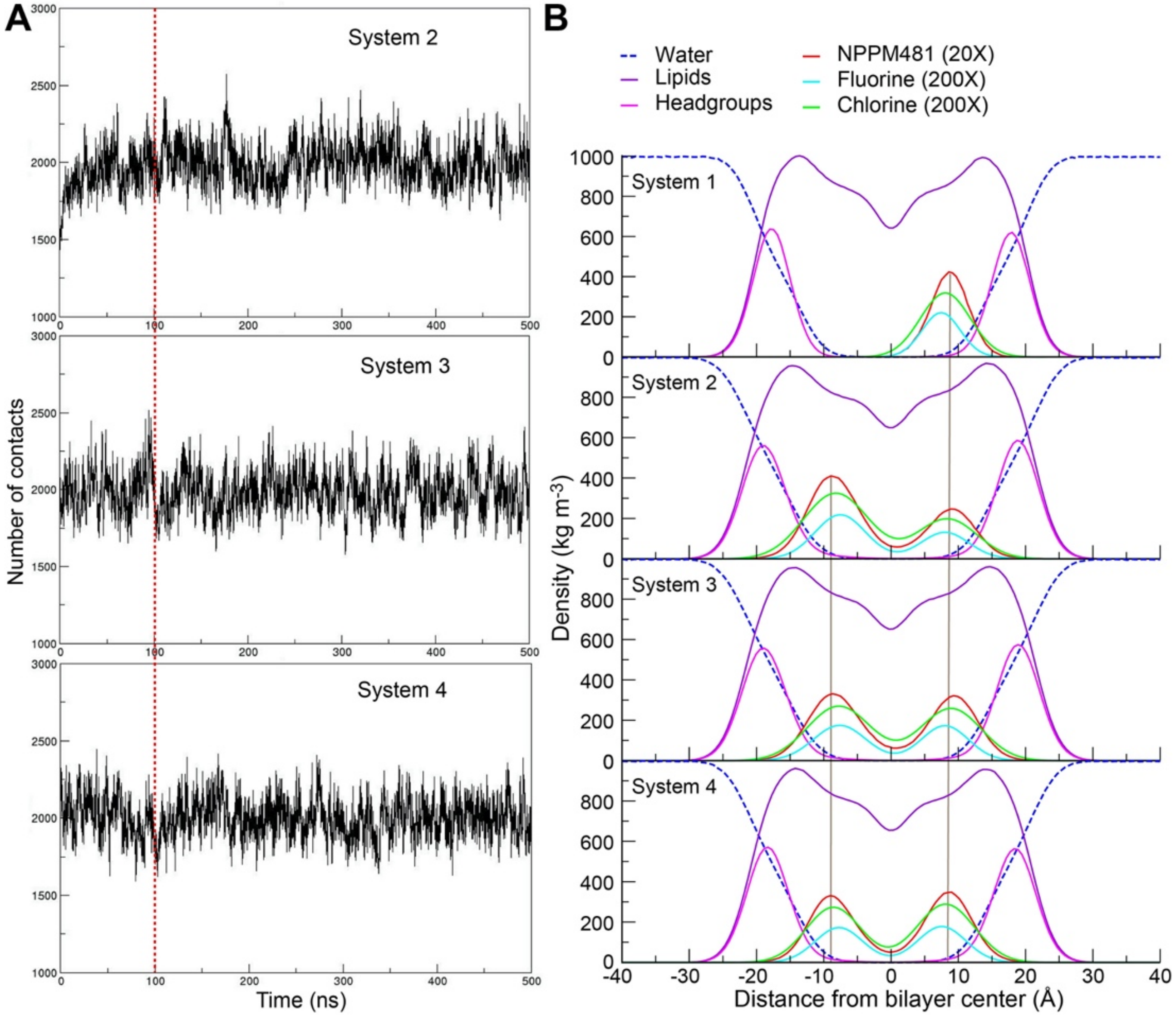


**Figure S8. Position of NPPM481 in membranes does not depend on initial simulation conditions.** The simulated systems are defined as follows: in system 1, one NPPM481 molecule was placed in solution 20 Å above the bilayer; in systems 2, 3 and 4, sixteen NPPM481 molecules were pre-inserted into the membrane at the bilayer center (system 2), the hydrocarbon region (system 3, eight molecules per leaflet), and the headgroup region (system 4, eight molecules per leaflet). The bilayer is made of 128 x 128 DMPC molecules. **(A)** The number of contacts between NPPM481 and the lipid headgroup atoms, plotted along the simulation trajectories for systems 2, 3, and 4. To ensure that the systems reached full equilibration, the first 100 ns (point marked with a red vertical dotted line) were not included in the subsequent analysis. **(B)** Mass density profiles calculated for the NPPM481 systems 1-4. Irrespective of the initial conditions, NPPM481 molecules and their Cl (A-ring) and F (B-ring) substituents converge to the same position when projected onto the membrane normal. The peak densities for the NPPM481 molecules are shown with vertical lines to guide the eye.

### TABLES

**Table S1.** X-ray data collection and refinement statistics. All complexes exhibit a 1:1 stoichiometry of SMI to Sec14. All crystal lattices are of the trigonal P3221 space group, and all contain one protein chain per asymmetric unit (AU) with exception of the himbacine complex which contains two. The two protein chains of the Sec14::himbacine complex in the AU adopt similar conformations with an overall root-mean-square-deviation (RMSD) = 0.4 Å. Parentheses indicate highest shell.

| **Data collection** | | | | | |
|---|---|---|---|---|---|
| Crystal | **NPPM481** (7ZGC) | **NPPM244** (7ZGD) | **NPBB112** (7ZGB) | **Ergoline** (7ZGA) | **Himbacine** (7ZG9) |
| Wavelength (Å) | 1.0000 | 1.0000 | 0.9999 | 0.9999 | 0.9999 |
| Space group | P3221 | P3221 | P3221 | P3221 | P3221 |
| a, b, c (Å) | 86.22, 86.22, 109.66 | 86.73, 86.73, 109.61 | 88.65, 88.65, 99.10 | 87.30, 87.30, 109.80 | 86.60, 86.60, 148.75 |
| $\alpha, \beta, \gamma$ (°) | 90, 90, 120 | 90, 90, 120 | 90, 90, 120 | 90, 90, 120 | 90, 90, 120 |
| Resolution (Å) | 50 – 2.24 (2.37-2.24) | 50 – 2.08 (2.21-2.08) | 50 – 2.70 (2.86-2.70) | 50 – 2.30 (2.44-2.30) | 50 – 1.76 (1.87-1.76) |
| $R_{meas}$ | 0.09 (1.49) | 0.11 (1.78) | 0.14 (1.95) | 0.09 (0.97) | 0.07 (1.55) |
| $I/\sigma I$ | 16.47 (1.42) | 14.52 (1.12) | 8.29 (0.82) | 13.02 (1.41) | 19.16 (1.25) |
| Completeness (%) | 99.7 (98.4) | 100.0 (99.9) | 99.5 (98.4) | 99.6 (98.0) | 100.0 (99.8) |
| Redundancy | 10.40 (9.88) | 10.40 (10.53) | 10.40 (9.88) | 4.54 (3.81) | 10.12 (9.44) |
| Unique reflections | 44052 (7032) | 55450 (8978) | 23922 (3836) | 41643 (6630) | 122833 (19843) |
| $CC_{1/2}$ | 0.999 (0.649) | 0.999 (0.502) | 0.996 (0.423) | 0.998 (0.593) | 1.000 (0.536) |

| **Refinement** | | | | | |
|---|---|---|---|---|---|
| *R-factor* | 0.1844 | 0.1976 | 0.2182 | 0.1883 | 0.1864 |
| $R_{free}$ | 0.2111 | 0.2316 | 0.2490 | 0.2284 | 0.2198 |
| Bond angles, rmsd (°) | 0.907 | 0.921 | 1.141 | 1.021 | 0.968 |
| Bond lengths, rmsd (Å) | 0.009 | 0.008 | 0.010 | 0.007 | 0.007 |

| **Ramachandran plot** | | | | | |
|---|---|---|---|---|---|
| Ramachandran favored (%) | 96.6 | 96.6 | 92.2 | 97.3 | 98.1 |
| Ramachandran allowed (%) | 3.1 | 3.4 | 7.5 | 2.7 | 1.9 |
| Ramachandran outliers (%) | 0.3 | 0 | 0.3 | 0 | 0 |

**Table S2.** NPPM481 atom types assigned by the CGENFF program.

| S.N. | Atom Name | Atom Types | S.N. | Atom Name | Atom Type |
|---|---|---|---|---|---|
| 1 | CL1 | CLGR1 | 21 | F1 | FGR1 |
| 2 | C2 | CG2R61 | 22 | C17 | CG2R6 |
| 3 | C3 | CG2R61 | 23 | C16 | CG2R6 |
| 4 | C4 | CG2R61 | 24 | C15 | CG2R6 |
| 5 | C1 | CG2R61 | 25 | C14 | CG2R6 |
| 6 | N1 | NG2O1 | 26 | H1 | HGR62 |
| 7 | O1 | OG2N1 | 27 | H2 | HGR61 |
| 8 | O2 | OG2N1 | 28 | H3 | HGR61 |
| 9 | C6 | CG2R61 | 29 | H4 | HGA2 |
| 10 | C5 | CG2R61 | 30 | H5 | HGA2 |
| 11 | C7 | CG2O1 | 31 | H6 | HGA2 |
| 12 | O3 | OG2D1 | 32 | H7 | HGA2 |
| 13 | N2 | NG2S0 | 33 | H8 | HGA2 |
| 14 | C8 | CG321 | 34 | H9 | HGA2 |
| 15 | C9 | CG321 | 35 | H10 | HGA2 |
| 16 | C10 | CG321 | 36 | H11 | HGA2 |
| 17 | C11 | CG321 | 37 | H12 | HGR61 |
| 18 | N3 | NG301 | 38 | H13 | HGR61 |
| 19 | C12 | CG2R61 | 39 | H14 | HGR61 |
| 20 | C13 | CG2R66 | 40 | H15 | HGR62 |
| | | | 41 | LP1 | LPH |

**Table S3.** QM and MM water interaction energies and distances in NPPM481.

| | Water interaction energy (kcal/mol) | | | Water interaction distance (Å) | | |
|---|---|---|---|---|---|---|
| Sites | QM | MM | Difference | QM | MM | Difference |
| C2 | -1.513 | -1.598 | -0.086 | 3.857 | 3.537 | -0.320 |
| C3 | -1.116 | -1.406 | -0.290 | 3.846 | 3.446 | -0.400 |
| C4 | -1.355 | -0.561 | 0.794 | 3.733 | 3.493 | -0.240 |
| C5 | -2.075 | -2.089 | -0.014 | 3.809 | 3.469 | -0.340 |
| C6 | -0.350 | -0.116 | 0.233 | 4.402 | 4.782 | 0.380 |
| C7 | 0.063 | -0.456 | 0.519 | 4.082 | 3.682 | -0.400 |
| C12 | -1.322 | -2.016 | -0.693 | 4.062 | 3.782 | -0.280 |
| C13 | -1.471 | -2.962 | -1.491 | 3.843 | 3.443 | -0.400 |
| C14 | -1.654 | -1.523 | 0.131 | 3.724 | 3.484 | -0.240 |
| C15 | -1.687 | -1.806 | -0.119 | 3.702 | 3.302 | -0.400 |
| C16 | -2.115 | -2.304 | -0.189 | 3.686 | 3.326 | -0.360 |
| C17 | -2.153 | -2.329 | -0.176 | 3.629 | 3.309 | -0.320 |
| F1 | -2.956 | -4.229 | -1.272 | 3.085 | 2.945 | -0.140 |
| N1 | -0.659 | -1.332 | -0.672 | 5.125 | 4.725 | -0.400 |
| N2 | -0.783 | -0.945 | -0.162 | 3.362 | 3.322 | -0.040 |
| N3 | -6.049 | -5.719 | 0.330 | 3.146 | 3.106 | -0.040 |
| O1 | -2.670 | -2.840 | -0.170 | 3.225 | 3.185 | -0.040 |
| O1 | -4.774 | -4.388 | 0.387 | 3.133 | 2.933 | -0.200 |
| O1 | -3.321 | -4.345 | -1.024 | 3.172 | 2.892 | -0.280 |
| O2 | -4.643 | -4.389 | 0.254 | 3.144 | 2.944 | -0.200 |
| O2 | -3.305 | -4.523 | -1.217 | 3.199 | 2.899 | -0.300 |
| O3 | -6.931 | -6.268 | 0.663 | 3.010 | 2.93 | -0.080 |
| O3 | -6.039 | -6.078 | -0.039 | 2.990 | 2.83 | -0.160 |
| LP1 | -1.310 | -1.395 | -0.085 | 3.192 | 3.212 | 0.020 |
| H1 | -3.446 | -2.855 | 0.590 | 2.352 | 2.492 | 0.140 |
| H2 | -3.281 | -2.080 | 1.200 | 2.269 | 2.649 | 0.380 |
| H3 | -4.164 | -3.830 | 0.334 | 2.293 | 2.593 | 0.300 |
| H4 | -2.128 | -1.958 | 0.170 | 2.569 | 2.729 | 0.160 |
| H5 | -0.198 | 0.496 | 0.694 | 2.604 | 2.964 | 0.360 |
| H6 | -2.419 | -1.792 | 0.627 | 2.516 | 2.776 | 0.260 |
| H7 | -2.086 | -1.503 | 0.583 | 2.580 | 2.78 | 0.200 |
| H9 | -2.702 | -2.428 | 0.274 | 2.477 | 2.677 | 0.200 |
| H10 | -1.875 | -1.855 | 0.020 | 2.752 | 2.792 | 0.040 |
| H11 | -0.332 | 0.754 | 1.086 | 3.332 | 3.712 | 0.380 |

| | | | | | | |
|---|---|---|---|---|---|---|
| H12 | -3.029 | -2.645 | 0.384 | 2.767 | 2.827 | 0.060 |
| H13 | -2.217 | -1.578 | 0.639 | 2.551 | 2.771 | 0.220 |
| H14 | -2.223 | -1.479 | 0.744 | 2.537 | 2.777 | 0.240 |
| H15 | -2.155 | -2.367 | -0.212 | 2.483 | 2.463 | -0.020 |
| CL1 | -0.757 | -1.399 | -0.642 | 3.977 | 3.577 | -0.400 |
| ***RMSE*** | | | ***0.617*** | | | ***0.267*** |

**Table S4.** Partial charges of NPPM481 atoms prior to and after optimization. Atoms for which partial charges were optimized and the corresponding charge values are shown in boldface.

| S.N. | Atom Name | Initial charge | Optimized charge | S.N. | Atom Name | Initial charge | Optimized charge |
|---|---|---|---|---|---|---|---|
| **1** | **CL1** | **-0.169** | **-0.166** | **21** | **F1** | **-0.223** | **-0.251** |
| **2** | **C2** | **0.151** | **0.255** | 22 | C17 | -0.201 | -0.201 |
| **3** | **C3** | **-0.212** | **-0.362** | 23 | C16 | -0.073 | -0.073 |
| **4** | **C4** | **-0.050** | **0.072** | 24 | C15 | -0.168 | -0.168 |
| **5** | **C1** | **0.296** | **0.109** | **25** | **C14** | **-0.076** | **-0.069** |
| 6 | N1 | 0.403 | 0.403 | 26 | H1 | 0.184 | 0.184 |
| 7 | O1 | -0.340 | -0.340 | 27 | H2 | 0.115 | 0.115 |
| 8 | O2 | -0.340 | -0.340 | 28 | H3 | 0.160 | 0.16 |
| **9** | **C6** | **-0.138** | **-0.096** | 29 | H4 | 0.090 | 0.09 |
| **10** | **C5** | **-0.187** | **0.069** | 30 | H5 | 0.090 | 0.09 |
| **11** | **C7** | **0.571** | **0.334** | 31 | H6 | 0.090 | 0.09 |
| **12** | **O3** | **-0.475** | **-0.475** | 32 | H7 | 0.090 | 0.09 |
| **13** | **N2** | **-0.785** | **-0.302** | 33 | H8 | 0.090 | 0.09 |
| **14** | **C8** | **0.121** | **0.028** | 34 | H9 | 0.090 | 0.09 |
| **15** | **C9** | **0.091** | **-0.144** | 35 | H10 | 0.090 | 0.09 |
| **16** | **C10** | **0.121** | **0.028** | 36 | H11 | 0.090 | 0.09 |
| **17** | **C11** | **0.091** | **-0.144** | 37 | H12 | 0.115 | 0.115 |
| **18** | **N3** | **-0.506** | **-0.217** | 38 | H13 | 0.115 | 0.115 |
| **19** | **C12** | **0.248** | **0.304** | 39 | H14 | 0.115 | 0.115 |
| **20** | **C13** | **0.093** | **-0.008** | 40 | H15 | 0.183 | 0.183 |
| | | | | 41 | LP1 | 0.050 | 0.050 |

**TableS5**. Dihedral angle parameters optimized using ffTK.

| S.N. | Atom type | Atom Type | Atom type | Atom Type | *K*φ[a] | n[b] | δ[c] |
|---|---|---|---|---|---|---|---|
| **1** | NG2O1 | CG2R61 | CG2R61 | CLGR1 | 6.9990 | 1 | 180.0 |
| | NG2O1 | CG2R61 | CG2R61 | CLGR1 | 2.0840 | 2 | 180.0 |
| **2** | NG301 | CG2R61 | CG2R66 | FGR1 | 6.5980 | 2 | 180.0 |
| **3** | NG2S0 | CG321 | CG321 | NG301 | 5.3440 | 1 | 180.0 |
| | NG2S0 | CG321 | CG321 | NG301 | 1.3090 | 3 | 0.0 |
| **4** | CG2R61 | CG2O1 | NG2S0 | CG321 | 1.6510 | 1 | 0.0 |
| | CG2R61 | CG2O1 | NG2S0 | CG321 | 2.8090 | 2 | 180.0 |

[a]*K*φ is the force constant for the dihedral angle term;
[b]n is the dihedral angle multiplicity;
[c]δ is the dihedral angle phase.
